# Readout of histone methylation by Trim24 locally restricts chromatin opening by p53

**DOI:** 10.1101/2022.08.23.504916

**Authors:** Luke Isbel, Murat Iskar, Sevi Durdu, Ralph S. Grand, Joscha Weiss, Eric Hietter-Pfeiffer, Zuzanna Kozicka, Alicia K. Michael, Lukas Burger, Nicolas H. Thomä, Dirk Schübeler

## Abstract

The genomic binding sites of the transcription factor (TF) and tumour suppressor p53 are unusually diverse in regards to their chromatin features, including histone modifications, opening the possibility that chromatin provides context-dependence for p53 regulation.

Here, we show that the ability of p53 to open chromatin and activate its target genes is indeed locally restricted by its cofactor Trim24. Trim24 binds to both p53 and unmethylated lysine 4 of histone H3, thereby preferentially locating to those p53 sites that reside in closed chromatin, while it is deterred from accessible chromatin by lysine 4 methylation.

The presence of Trim24 increases cell viability upon stress and enables p53 to impact gene expression as a function of the local chromatin state.

These findings link histone methylation to p53 function and illustrate how specificity in chromatin can be achieved, not by TF-intrinsic sensitivity to histone modifications, but by employing chromatin-sensitive cofactors which locally modulate TF function.

## Introduction

Transcription factors (TFs) are DNA-binding proteins that determine distinct spatial and temporal transcriptional patterns. Beyond DNA sequence, chromatin proteins such as nucleosomes and their modifications are thought to add an additional layer of in gene regulation^1-3^. However, it remains largely unclear how these modifications are ‘read-out’ by TFs, as these lack known domains that specifically interact with histones. However, such domains are abundant on non-DNA binding transcriptional cofactors^4,5^.

The base unit of chromatin is a nucleosome, that consists of ∼147bp of DNA wrapped around a histone octamer. This arrangement sterically hinders access to 95% of the normally exposed DNA binding surface for TF engagement, making nucleosomes intrinsically antagonistic to TFs^6,7^. Nucleosome modifications such as histone H3 methylation of lysine 4 (H3K4me) or acetylation of lysine 27 (H3K27ac) play a functional role in the actions of TFs, but the mechanism by which they do so remains enigmatic^2,8^. Modifications might directly affect TF access to DNA by altering nucleosome-DNA wrapping kinetics^9^. Alternatively, TFs may recruit cofactors that are sensitive to histone modifications. For example, many chromatin remodeler complexes possess a combination of chromatin-binding domains that modulate their activity, at least *in vitro*^8,10,11^. In the latter case, it seems possible that such cofactors enable TF specificity to regulate gene activity on the basis of local chromatin states.

High accessibility and the presence of euchromatic histone modifications are hallmarks of active regulatory regions, such as enhancers and promoters, and can be successfully used for their detection^12,13^. This coincides with the binding of TFs, suggesting a functional link^14^. However, these striking correlations with chromatin features do not inform if a TF creates open chromatin or binds as a consequence thereof. Loss-of-function studies suggest that some TFs are at least partially involved in maintaining an ‘open’ chromatin state at regulatory regions^15^. This makes it highly challenging to disentangle the contribution of ‘naïve’ chromatin towards the initial engagement of a TF, from the ongoing remodelling that occurs due to TF binding at a steady state^16^. These issues are partially circumvented by studying TF binding upon induction or ectopic expression, as this enables to monitor the initial TF engagement and should reveal the sensitivity to chromatin^16^. Of particular interest has been the binding to regions of closed chromatin by so-called ‘pioneer TFs’ to drive cell-fate and differentiation^17^. *In vitro*, several TFs have been demonstrated to engage nucleosomal DNA, though it remains to be determined if this behaviour occurs *in vivo* and if histone modifications alter nucleosomal engagement^18,19^.

The TF and tumour suppressor p53 controls the expression of numerous genes involved in DNA repair, cell cycle arrest and cell death^20^. p53 can be rapidly induced in response to various forms of cellular stress, resulting in immediate activation of target genes^21,22^. p53 is unusual in that only a fraction of p53-bound sites show signs of open chromatin under basal conditions^22-25^. Thus, p53 seems to engage closed chromatin sites without creating accessibility until activated by stress-response pathways. However, it remains unknown how stable engagement in closed chromatin is achieved or indeed if there are functional consequences from such binding.

Here, we show that a cofactor, Trim24, limits the degree to which p53 can create accessibility in closed chromatin and furthermore, that it does so by reading out the local methylation state of lysine 4 of histone 3. This provides a molecular link between histone methylation and p53 function, enabling locus specific TF regulation subsequent of its initial binding.

## Results

### p53 engages closed chromatin in the mouse and human genome

Features of open chromatin such as accessibility, ‘active’ histone marks and low DNA methylation are hallmarks of sites of TF binding^14,26-28^. p53 appears different in this respect as it has been reported to occupy sites of closed and open chromatin^22-24^. We investigated p53 binding in mouse embryonic stem cells (mESCs), which represent a non-transformed cell type. Under ‘basal’ cellular conditions, p53 displayed strong binding at both open and closed chromatin loci, with cellular stress-induced activation of p53 via doxorubicin treatment resulting in increased binding and accessibility at these sites (Fig. 1a and Supplementary Fig. 1a-h)^29,30^. This diversity of chromatin states at p53 binding sites is also evident in human tissues and ESCs (Fig. 1b and Supplementary Fig. 2a-f) and is in stark contrast to TFs previously shown to be chromatin insensitive, for which binding almost exclusively occurs in open chromatin and suggesting it is more typical for TFs to create accessibility upon closed chromatin binding (Supplementary Fig. 1h)^31^.

**Fig. 1.**
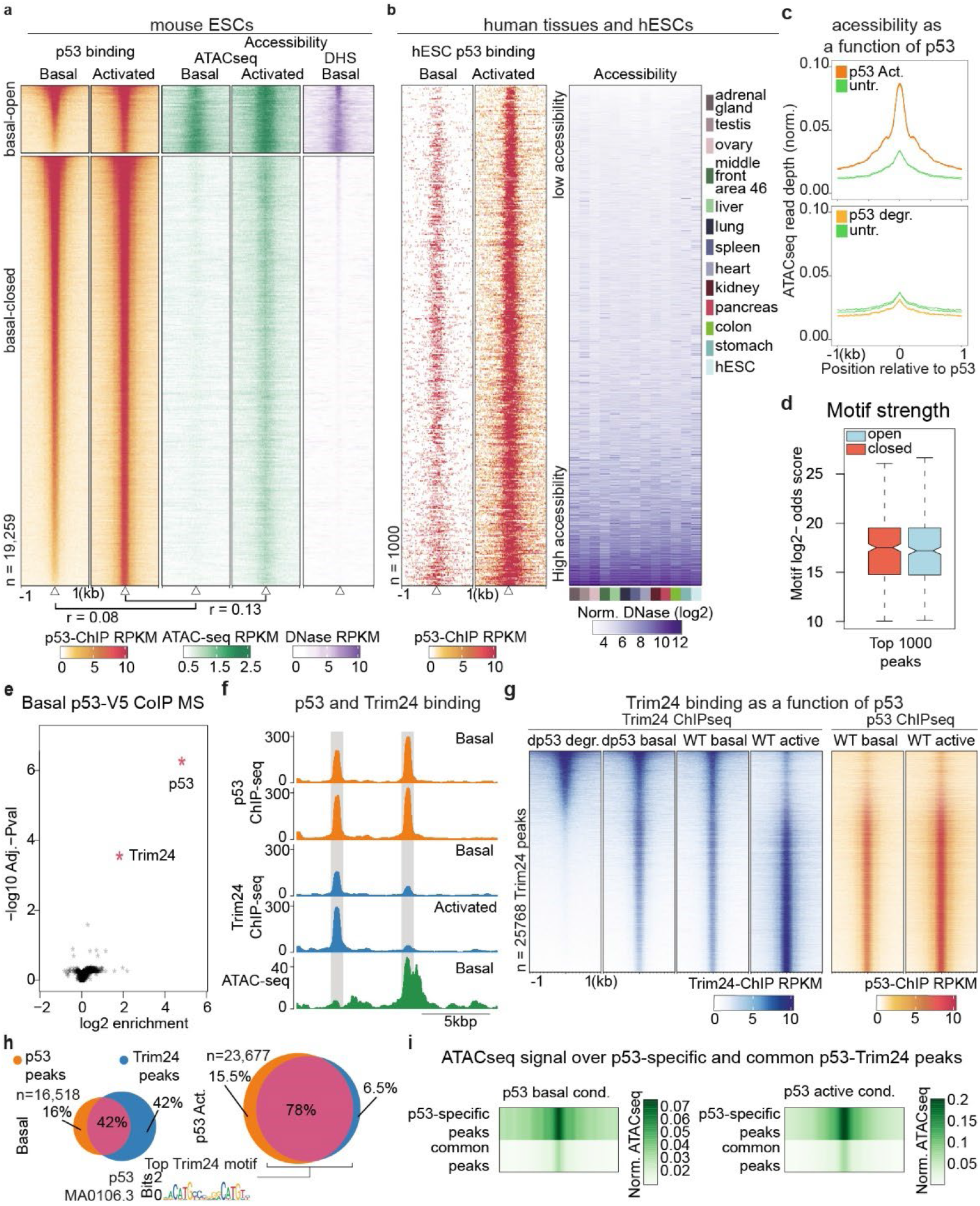
p53 binds closed chromatin in mESCs and human tissues and recruits Trim24. **a**) p53 binds to open and closed mESC chromatin. Binding and accessibility (ATACseq-middle, DNaseSeq-right) increase following activation (4hrs, doxorubicin-1µM), Pearson correlation score indicated (below). **b**) hESC-p53 ChIPseq (left) alongside DNaseSeq accessibility in human tissues and hESCs (right). **c**) Accessibility metaprofile (normalized ATACseq signal) in WT (top) and degron-tagged p53 (bottom) cells after activation and degradation (dTAG13 500nM, 4hrs), respectively. **d**) Motif score (log2-odds) at open and closed chromatin sites. **e**) CoIP mass spectrometry (MS) of V5-tagged p53 enriches for both p53 and Trim24 proteins, indicated in red are proteins with adj. *P* value < 0.01. **f**) p53 and Trim24 binding (ChIPseq) and accessibility (ATACseq), highlighted are representative p53-Trim24 co-bound and p53 only sites. **g**) Heatmaps of Trim24 ChIPseq upon p53 degradation in the dp53 degron tagged line and upon activation in wildtype mESCs (WT) (left), alongside p53 ChIPseq in WT cells (right). **h**) The overlap in p53 and Trim24 peaks under p53-basal (left) and -active (right) conditions. The top motif from Trim24 ChIPseq peaks is shown. **i**) ATACseq in basal (left) and active (right) conditions at common and p53-only peaks as in **h**. p53-specific peaks overlap accessible regions, while common Trim24-bound peaks are relatively depleted in accessibility.

**Fig. 2.**
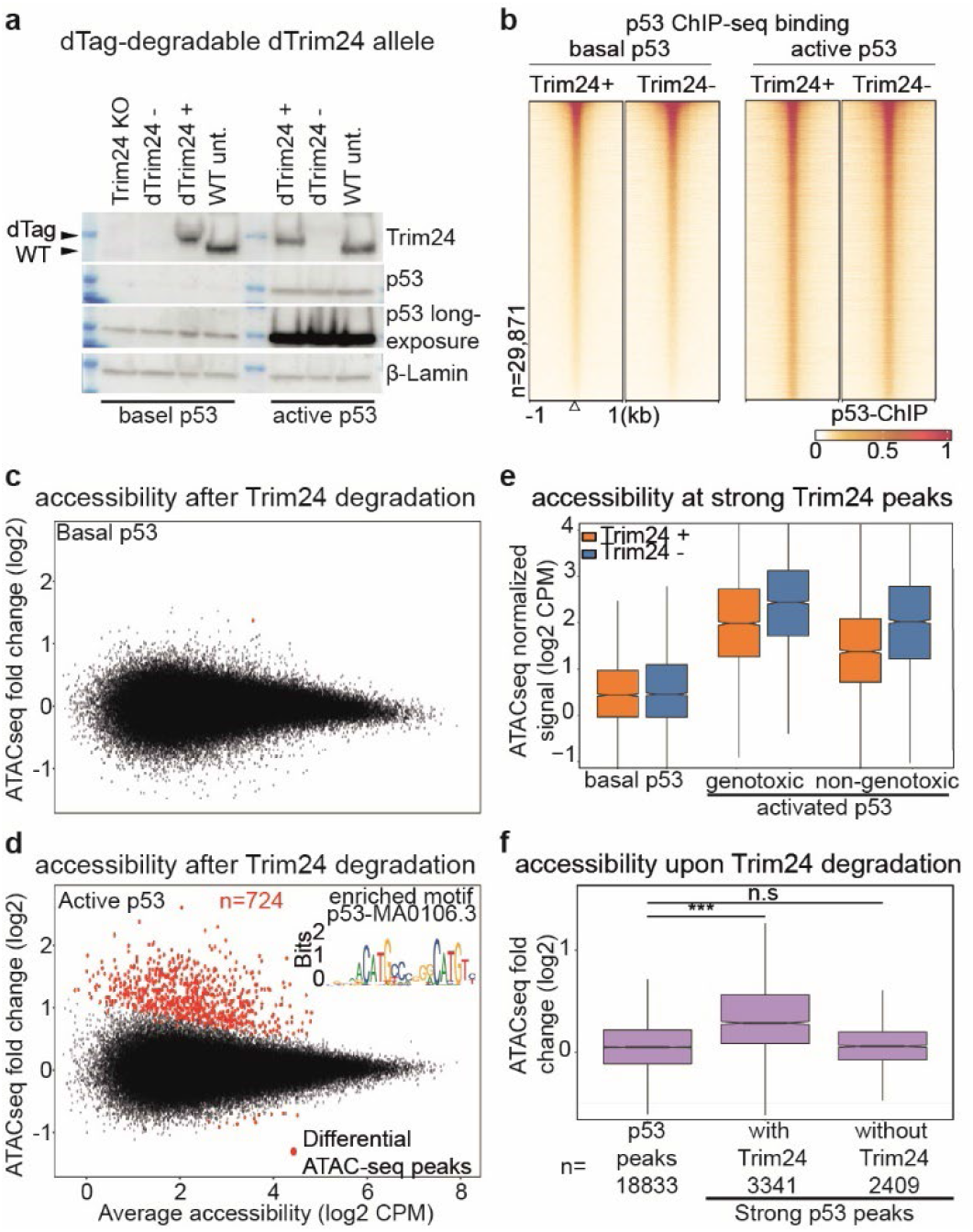
Trim24 is a negative regulator of active p53 in the genome. **a**) Tagged Trim24 can be rapidly degraded including during p53 activation (dTAG13 500nM, Doxorubicin 1uM – 4hrs). **b**) p53 ChIPseq in Trim24-degraded mESCs at combined p53 and Trim24 peaks. **c** & **d**) MA plot of changes in accessibility (log2 CPM) upon degradation of Trim24, under basal (top) and active (bottom) conditions, for all mESC ATACseq peaks (n∼165K). p53 motifs are enriched at differential sites in active conditions (*P* value < 1e-297). **e**) ATACseq signal (log2 CPM) under basal, doxorubicin (genotoxic) and nutlin3a (non-genotoxic) p53-activated conditions for 4 hours, with and without Trim24 loss at strong Trim24 peaks (n= 3883), i.e., > 3.5-fold log2 Trim24 ChIPseq enrichment. **f**) Increase in ATACseq signal at all p53 ChIPseq peaks and strong p53 peaks with or without Trim24, i.e. > 3.5-fold log2 ChIPseq enrichment for p53 and/or Trim24.

Several reports suggest p53 may be responsible for enabling repressive chromatin marks at binding sites^32^. To test if p53 influences chromatin states under basal conditions, we tagged it to enable acute and targeted degradation. Specifically, we integrated V5 and dTAG peptides at the endogenous gene, allowing detection by the V5 epitope and dTAG-inducible degradation (Supplementary Fig. 3a-b)^33^. This ‘*dp53’* allele displayed similar expression and genomic binding compared to WT p53 under basal conditions (Supplementary Fig. 3a-b). Upon stress-activation it retains functionality, though to a lesser degree, as measured by binding and the ability to create accessible chromatin (Supplementary Fig. 3c-m). Rapid degradation of p53 resulted in a slight but significant decrease in accessibility at open chromatin, but no increase at closed chromatin, arguing against a role for p53 in its maintenance (Fig. 1c and Supplementary Fig. 3j-k). These results suggest that chromatin is largely agnostic to p53 prior to activation and opens the possibility that features of closed chromatin enable stable binding of p53.

**Fig. 3.**
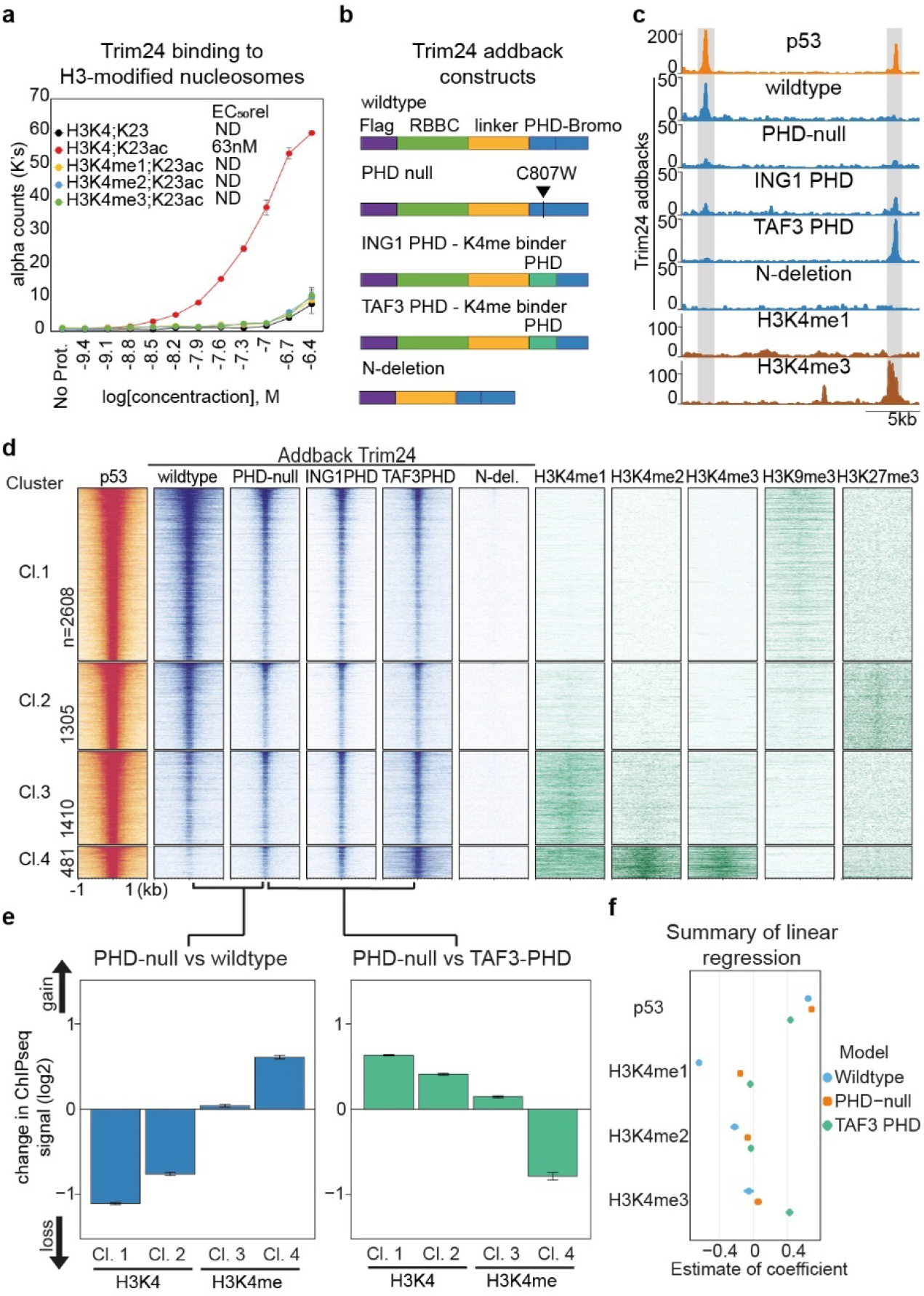
Trim24 impacts p53 in a H3K4 methylation-dependent manner. **a**) Binding of recombinant Trim24 to synthetic modified nucleosomes. **b**) Constitutively expressed addback alleles in a *Trim24* KO background, including mutations that either nullify the PHD domain or invert its preferences towards methylated H3K4. **c**) Representative regions (grey) of p53- and addback Trim24-binding in open and closed chromatin. **d**) K-means clustering of strong p53-binding sites (n=5804 sites) by histone modifications. Shown are histone marks representative of each cluster (right), alongside binding of p53 and Trim24 variants (left). **e**) Change in binding of the PHD-null Trim24 relative to wildtype (left) and TAF3-PHD variants, for each of the clusters shown in d. Displayed are the mean fold change, error bars indicate std. err. **f**) Estimate of coefficient contributions to linear regression model of binding for wildtype, PHD-null and TAF3-PHD variants, using p53 and H3K4 methylation enrichment as variables.

### p53 binding in closed chromatin is independent of DNA methylation

One feature of closed chromatin is high levels of DNA methylation, which could be potentially relevant for p53 as it has been reported to prefer some motif variants when methylated^25^. We therefore asked if DNA methylation could account for the observed binding patterns. If true, we would expect a reduction in binding at closed chromatin sites upon removal of DNA methylation. To test this, we measured p53 binding by ChIPseq in cells that lack DNA methylation due to genetic deletion of all three DNA methyltransferases – DNMT-TKOs^34^. In these cells p53 binding was largely indistinguishable from WT, with continued binding in closed chromatin (Supplementary Fig. 4a-b). Similarly, comparing the strongest sites located in either closed or open chromatin revealed similar motif strength (Fig. 1d). We conclude that the binding of p53 to closed chromatin appears independent of DNA methylation.

**Fig. 4.**
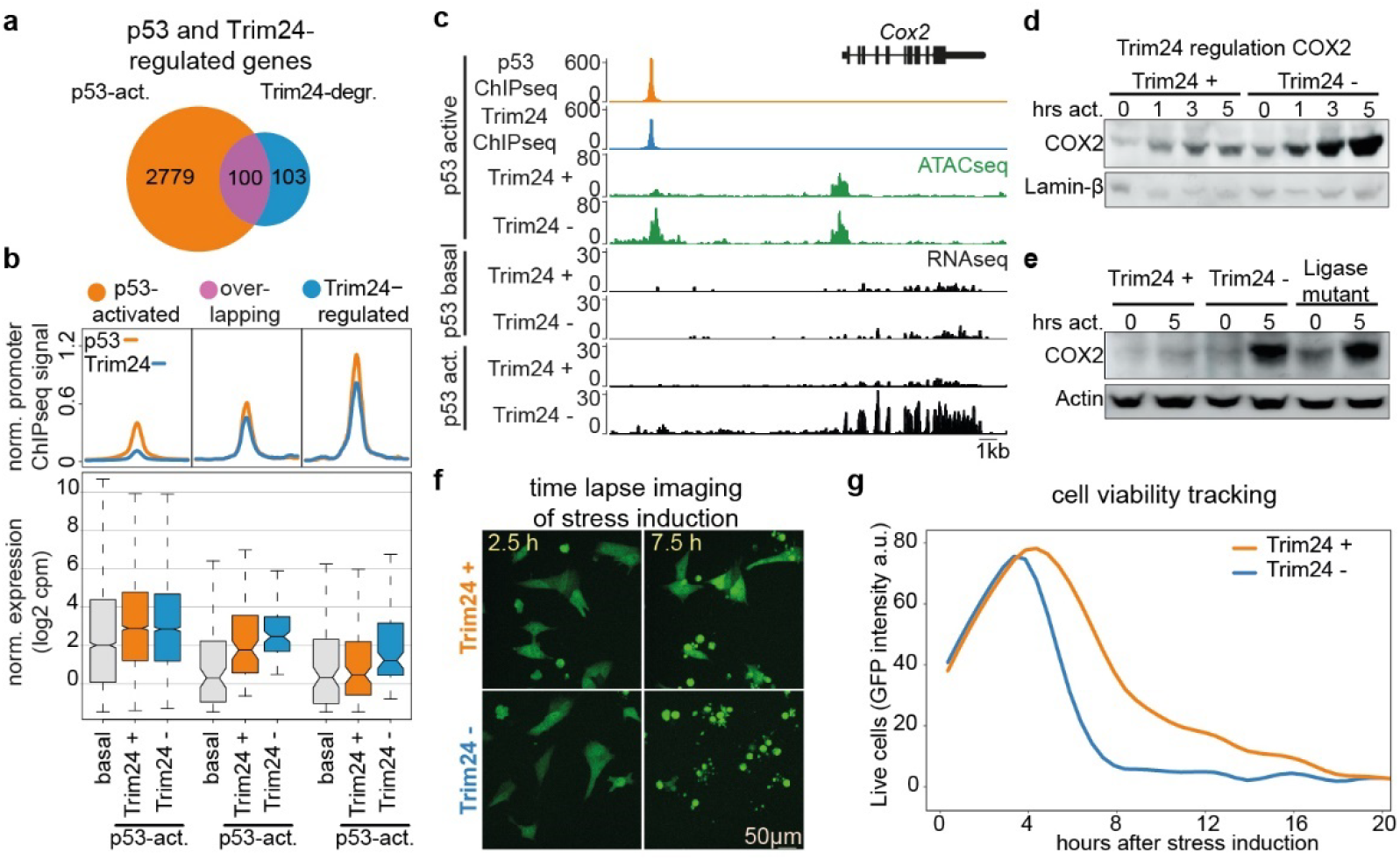
Trim24 regulates a subset of p53-target genes. **a**) Number and overlap of genes that change upon p53-activation compared to basal cells and upon Trim24 degradation in p53-active conditions. **b**) The p53 and Trim24 ChIPseq signal (top) in promoter peaks of deferentially expressed genes as in **a**, alongside gene expression (bottom) in basal and p53-activated conditions with and without Trim24. **c**) Representative Trim24 regulated gene, *Cox2*. Shown is p53 and Trim24 binding and accessibility, and RNAseq tracks under either basal or p53-activated conditions and with or without Trim24 loss. **d**) Western blot of COX2 protein expression after p53-activation, with and without Trim24. **e**) Western blot of COX2 protein expression after p53-activation, with and without Trim24 and in the ligase-null mutant (C52/55A) Trim24 variant line. **f**,**g**) Live imaging of GFP expressing mESCs, measuring cell viability across a 20-hour period following stress induction and with or without Trim24-degradation. **f**) representative images at 2.5 and 7.5 hours post stress induction. **g**) Cell viability as a function of Trim24, quantified for 20 hours following stress induction, a.u. arbitrary units.

### Trim24 locates with p53 in the genome

We speculated that binding patterns could be explained by p53 cofactors that are able to bind to chromatin. To identify potential cofactors, we carried out CoIP-MS from whole cell extract using the V5 tag on the dp53 allele. Encouragingly, p53 itself was the top enriched protein (Fig. 1e and Supplementary Fig. 5a-b), followed by the Trim24 protein with comparable enrichment, which is a protein that possesses canonical histone binding domains. Trim24 has previously been implicated as a p53-interacting E3-ubiquitin ligase^35^. Trim24 has no DNA binding ability on its own, yet it contains PHD- and bromo-domains that can interact with modified histones^36^, opening the possibility that Trim24 links p53 binding to local chromatin.

To determine if the observed interaction translates into co-occupancy across the genome, we carried out ChIPseq of Trim24 under both basal and p53-active conditions (Fig. 1f and Supplementary Fig. 5c-e). The top enriched sequence at Trim24 peaks was the p53 motif and we detected co-occupancy at the majority of peaks, with a high degree of scaling in the signal for both factors. To ask if Trim24 depends on p53 for binding, we measured Trim24 binding upon p53 degradation using the p53 degron line (Fig. 1g and Supplementary Fig. 5f). Loss of p53 leads to absence of Trim24 at p53 sites, suggesting that p53 recruits Trim24. In addition, we monitored binding upon p53-activation, which coincided with an increase in Trim24 binding at p53 sites. These data argue that Trim24 binding corresponds to both the presence and levels of p53. While true for the majority of genomic sites, we noticed a subgroup bound exclusively and strongly by p53 (Fig. 1f,h and Supplementary Fig. 5e). Comparing the ATACseq signal at overlapping against non-overlapping Trim24 and p53 peaks revealed that co-occupied sites showed low to no accessibility and thus reside in closed chromatin (Fig. 1i). In contrast, sites bound by p53 alone were enriched in accessibility and reside in open chromatin. Together, these data raise the possibility that the binding of Trim24 at p53 sites depends not only on p53 but also on the local chromatin state.

### Trim24 is not required for p53 binding to closed chromatin

To test if Trim24 contributes to p53 genomic binding we endogenously tagged *Trim24* with the V5 and dTAG peptides that enable detection and acute degradation (Fig. 2a). Trim24 degradation occurred rapidly but did not affect expression levels of the p53 protein or decrease its genomic binding (Fig. 2a,b and Supplementary Fig. 5g-h). We retested this at cellular resolution using immunofluorescence microscopy, which confirmed the minimal effects after Trim24 removal (Supplementary Fig. 6a-e). This was true for basal as well as active p53-conditions. Indeed, only a small number of p53 sites (n∼150) appear to respond to the absence of Trim24 by showing increased p53 binding (Supplementary Fig. 5h).

### Trim24 represses p53-dependent accessibility in active conditions

As Trim24 was not required for binding, we asked if it were recruited to modulate the ability of p53 to remodel chromatin. We therefore profiled accessibility in cells upon rapid loss of Trim24 in p53-basal conditions (Fig. 2c and Supplementary Fig. 7a-b). This revealed almost no changes in accessibility genome-wide, in stark contrast to changes observed in p53-active conditions, where several hundred loci showed increased accessibility upon loss of Trim24 (Fig. 2d). These loci were highly enriched for p53 binding (Supplementary Fig. 7c-d). Only a few loci decreased in accessibility (n=11). To determine if the observed Trim24 function is specific to the cellular stress used to activate p53 and thus potentially limited to the utilized doxorubicin-induced DNA damage, we also exposed cells to two additional activation paradigms: MDM2 inhibition using nutlin3a and DNA damaging via ionizing radiation (IR) (Fig. 2e & Supplementary Fig. 7e-g)^37,38^. Regardless of the method used to activate p53, Trim24 loss increased accessibility at a set of strong p53 sites that respond to doxorubicin. This argues that Trim24 limits the ability of active-p53 to generate local chromatin accessibility.

### Trim24 acts on chromatin in a locus specific manner

Next, we asked if Trim24 affected all p53 sites, or if it was limited to only the closed chromatin subset of p53 sites where both bind (Fig. 1h). We therefore reanalysed accessibility at strong p53 peaks with and without Trim24 binding (Fig. 2f). This revealed an increase in accessibility at co-occupied p53-Trim24 peaks, but not at the p53-only peaks. Furthermore, these Trim24-dependent effects were present up to twelve hours after p53 activation (Supplementary Fig. 8a-c). Together, these data argue that the activity of Trim24 takes place at local genomic elements with specific chromatin features rather than generally within the nucleoplasm, in contrast to other direct regulators such as Mdm2 that act on total cellular p53 levels.

### Trim24 localization is modulated by H3K4 methylation

Thus, we tested the interaction specificity of the histone-binding domains of recombinant full-length mouse Trim24 towards a panel of modified nucleosomes via an amplified luminescence proximity homogeneous assay. Consistent with previous reports on the isolated PHD/Bromo-domain and histone tail-peptides, we detected high enrichment of H3K23ac-modified nucleosome, but only when H3K4 was not methylated (Fig. 3a & Supplementary Fig. 9a)^36^. In this *in vitro* assay Trim24 only demonstrated binding when both preferred substrates for the PHD- (H3K4 unmethylated) and Bromo-domain (H3K23 acetylated) were present, suggesting that these marks are required for histone engagement.

However, in the cellular context, Trim24 bound sites show no enriched signal for H3K23ac, or any additional active histone marks (Supplementary Fig. 9b-d). This is different in the case of unmethylated H3K4, which is indeed highly prominent at Trim24-bound sites (Supplementary Fig. 9b,d). This opens the possibility that the PHD domain of Trim24 could contribute to its genomic binding at p53 sites. We tested this hypothesis with a set of Trim24 mutants, which we ectopically expressed in a *Trim24* knockout background. Tested variants included a PHD-null point mutant^36^ and PHD-domain substitutions from ING1 and TAF3 proteins that bind only methylated H3K4^39-41^ (Fig. 3b-c and Supplementary Fig. 10a-d), as well as the wildtype (WT) Trim24 as a control. In this complementation assay WT Trim24 showed genomic binding comparable to that of endogenous Trim24, albeit with a slightly reduced signal, suggesting that p53 and chromatin binding preferences were recapitulated (Supplementary Fig. 10e). The N-terminal deletion variant did not bind p53 sites, arguing this part of Trim24 as the p53-interaction point.

Next, we asked how PHD domain variants influence Trim24 binding as a function of histone modifications. A single point mutation in the PHD domain that blocks the H3K4 interaction resulted in decreased binding in closed chromatin and increased binding in open chromatin, both at the single locus level and globally at p53 sites (Fig. 3c-f and Supplementary Fig. 10f). While this argues that the PHD domain is required for chromatin sensitivity, it further suggests that the specificity of the PHD domain is necessary for modulating binding at p53 sites. To test this, we examined binding of Trim24 with heterologous PHD domains possessing different specificity, that of ING1 reported to bind to HK4me2 and HK4me3 and that of TAF3 reported to preferentially bind HK4me3^39-41^. Indeed, if we embed these two domains in the Trim24 protein they are sufficient to redirect Trim24. Importantly, this happens in a chromatin-sensitive manner, leading to increased binding to regions of H3K4 methylation, thus inverting the preference of Trim24 for unmethylated HK4 (Fig. 3d-f). In summary, the PHD domain is necessary and sufficient for the observed chromatin sensitivity of Trim24.

We were keen to examine if Trim24 binding and impact on p53 is modulated by other chromatin marks characteristic for heterochromatin. More specifically we focussed on H3K9me3, involved in repeat silencing, and H3K27me3, a critical component of repression by the Polycomb system^42,43^. This revealed that p53 is able to bind regions with either of these two features, at least in mESCs (Supplementary Fig. 11a-d). Profiling ATACseq signal upon activation demonstrates that sites enriched in these marks respond to p53 activation and Trim24 degradation with increased accessibility (Supplementary Fig. 11e-f). Together, these data argue that H3K4 methylation readout by the PHD domain fully accounts for Trim24 sensitivity to open chromatin and sequesters it at closed chromatin sites where it modulates p53-function.

### Trim24 impacts the transcriptional response of p53-activation

Since Trim24 modulates p53-dependent accessibility, we asked if it contributes to p53-target gene regulation in the Trim24 dTag line. We profiled expression upon p53 activation in the absence or presence of Trim24 (Fig. 4a and Supplementary Fig. 12a). Trim24 loss resulted in 203 genes responding differently to p53 activation (Supplementary Fig. 12b-g). These are bound by p53 and Trim24 at promoter sites (Fig. 4b). Half of these genes were identified as p53-responding genes upon stress induction in the presence of Trim24 and showed an enhanced response in the absence of Trim24. The other half shows only a very weak p53 response that is specifically enhanced in the absence of Trim24. We conclude that Trim24 responsive genes at the transcriptional level are a defined subset of p53 target genes, in line with our observation of Trim24 binding and changes in accessibility. Importantly, Trim24 targets are transcriptionally inactive in p53-basal conditions and tend to respond weakly or not at all to activation, supporting a model whereby Trim24 represses p53-tagets in closed chromatin.

Given that Trim24 has previously been linked to the repression of repetitive DNA elements we profiled annotated mouse repeats which identifies six subtypes that increase in expression upon Trim24 removal (Supplementary Fig. 12h)^44^. This included RLTR1B, RLTR1F and MMGLN elements that possess p53 binding sites in their LTR-promoter sequences (Supplementary Fig. 12i). We note that these only represent a minority of repeats previously implicated to be repressed by p53^45-47^. Trim24-repressed genes include *Cyclooxygenase 2* – *Cox2*, which is a key therapeutic target of anti-inflammatory drugs and is known to be silent unless induced by inflammatory stimuli (Fig. 4c-d)^48,49^. Cox2 response to p53 activation in stem cells is strongly repressed by Trim24, as in its absence there is an at least nine-fold increase in expression both at the RNA and protein level.

### The RING domain is required for Trim24 function at p53 sites

The N-terminal RING domain is a characteristic feature of TRIM proteins, which in many cases confers E3 ubiquitin ligase activity^50^. At the same time, it has been difficult to identify substrates and to separate enzymatic activity of TRIM proteins from their potential scaffolding function, as is most evident for the highly studied Trim28^51^, which recruits Setdb1 that catalyses repressive H3K9me3 for silencing^52^. To determine the role of the RING domain in Trim24, we generated a specific point mutation at the endogenous gene using CRISPR/Cas9. This should result in a ubiquitin ligase-null mutant version of the protein, as previously shown for the homologous Trim33 protein^53^. Next, we determined the impact of this RING domain-mutant on p53 dependent gene activation, which revealed that it failed to efficiently silence Trim24-repressed genes, arguing for the requirement of its ligase activity for Trim24 function (Fig. 4e and Supplementary Fig. 13a-d).

### Trim24 contributes to cell viability upon stress

To examine if Trim24 target genes contribute to the phenotypic stress response in mESCs, we utilized live-cell tracking of proliferation and apoptotic events over a 40-hour time course of p53 activation and upon loss of Trim24. Absence of Trim24 resulted in a rapid decrease of live cells, while Trim24-expressing cells persisted for twice as long before reaching similar levels (Fig. 4f-g, Supplementary Fig. 14a and Supplementary movie 1). In p53 basal conditions, no change in cell death was detected, but loss of Trim24 slightly decreased proliferation (Supplementary Fig. 14b and Supplementary movie 2). Together, these data argue that Trim24 limits the activation potential of a subset of initially silent p53 target genes and is required for a normal stress response in cells.

## Discussion

Here we demonstrate that Trim24 bridges p53 activity to the local chromatin state via its interaction with p53 and unmethylated lysine 4 of histone H3. p53 has the unusual characteristic to bind its motifs embedded in heterochromatin, yet its ability to open chromatin and activate genes upon stimulation is impacted by local histone modifications in a Trim24-dependent process. In this context, it is important to note that almost all TFs do not possess known protein domains that recognize histone modifications, such as PHD, bromo, Chromo-domains etc., arguing that TFs generally do not directly interact with chromatin marks^5^. Our work provides an example of how a specific cofactor, Trim24, connects a TF with histone modifications. The activity of p53 and its target genes are tightly controlled and mediating such context-dependent activation might be particularly relevant for TFs that can engage heterochromatin. In the case of Trim24, this could serve as a mechanism to locally threshold p53 response or to enable cell type specific responses that vary with changes in lysine 4 methylation, which are well described e.g. for cell type specific enhancers^54^. Trim24 has previously been linked to cancer, showing elevated expression in several types including breast cancer, while genetic loss and gain of expression in mice leads to elevated rates of cancer^36,55-61^. This has motivated the development of Trim24 degrader compounds^62^. It is tempting to speculate that Trim24 might have a specific response on p53 activity in cancer, given the findings here demonstrating the mutual contribution of H3K4 methylation and p53 on gene expression.

We further identify *Cox2* as a prominent Trim24-regulated gene (Fig. 4c). It encodes a highly clinically relevant enzyme that catalyses the first, rate-limiting step of prostanoid formation and is the target for non-steroidal anti-inflammatory COX2 inhibitors (Coxibs), indicating a role in pain, fever, inflammation, and tumorigenesis^48,63-65^. A key theme of *Cox2* function is its status as an inducible gene activated upon inflammatory stimuli, in keeping with the role of Trim24 in repressing p53-response genes that are initially inactive (Fig. 4b). Its status as a Trim24 target suggests the need to tightly modulate *Cox2* activation and indeed, overexpression or prolonged use of COX-2 selective inhibitors results in carcinogenesis and cardiotoxicity^66,67^. As loss of Trim24 decreases cell viability upon stress, this opens the possibility that regulating the dosage of p53 targets such as *Cox2* is required for a normal stress response.

While p53 was responsible for the majority of Trim24-binding sites in mESCs, there are several reports of additional Trim24-interacting TFs in various cellular contexts^60,68,69^. Whether these additional interactions are sensitive to closed chromatin remains to be determined, though the potential for this is likely higher during cellular transitions such as differentiation, when newly active TFs initially encounter closed chromatin. Trim24 has been reported to have a role in the silencing of repetitive-elements in hepatocytes^44^, similar to its paralogs Trim28 and Trim33 that are recruited by TFs to silence specific repetitive elements^70,71^. Here, we observed a specific set of repeats with strong p53 motifs which become activated in mouse stem cells after Trim24 loss, arguing for a highly targeted repression system that is TF- and context-dependent. Trim24 is present at target loci but appears to only function upon p53 activation.

Trim24 loss did not alter p53 occupancy in closed chromatin and the nature of stable p53 binding at these regions remains to be determined (Fig. 2b). While other studies have reported that repressive chromatin features such as H3K9me3 are inherently restrictive for TF binding^72^, several high affinity p53-binding sites co-occupy these marks and these are made accessible by p53 activation (Supplementary Fig. 11a-f). Our results argue that, in the case of p53, specific chromatin marks can limit the ability of a TF to open chromatin, thereby restricting TF functions in a manner distinct from simply blocking DNA binding.

It remains to be determined how Trim24 exerts a repressive function on p53, as it does not appear to impact local p53 levels. Trim24 has previously been implicated as a ubiquitin ligase of p53^35^ and its RING domain is required in human cancer cell line models^62^. It has proven challenging to characterize E3 ligases substrates at endogenous protein levels^73,74^ or separate ligase activity from scaffolding function as in the case of Trim28^70^. Here, we show that Trim24 requires a functional RING domain to regulate p53 (Fig. 4e and Supplementary Fig. 13a-d). Trim24 has recently been demonstrated to enable K63-linked ubiquitination, and while K48-linked ubiquitination is associated with degradation, K63-linked ubiquitination is instead associated with modulating protein-protein interactions^75,76^. We were able to recapitulate Trim24-dependent effects upon activation of p53 via inhibition of Mdm2, which catalyses K48-linked ubiquitination (Supplementary Fig. 7g), arguing that Mdm2- and Trim24-dependent ligase activities on p53 are functionally distinct. It remains to be determined how Trim24 ligase activity modulates p53 activity or how such post-translation modifications influences p53-interactions with any of the several coactivator proteins previously described^77-81^.

Our findings reveal a mechanism by which TF-regulation is accomplished locally on chromatin and represent a convincing instance where a histone mark is functionally linked to TF potency by the actions of a single histone-interaction domain. It seems likely that other chromatin interacting proteins with histone-interacting domains could play comparable roles, which should be testable using a combination of high-resolution readout and precise functional models that we have applied here.

## Methods

### Data reporting

No statistical methods were used to predetermine sample size. The experiments were not randomized, and the investigators were not blinded to allocation during experiments and outcome assessment.

### Cell culture Mouse ES cells

Mouse ES cells (TC-1 line, background 129S6/SvEvTac, originally obtained from A. Dean at the National Institutes of Health) with an recombinase mediated cassette exchange (RMCE) site located in the gamma-globin locus were used as wildtype cells^82^. These were used for the generation of p53 and Trim24 degron lines. Mouse ES cells were cultured as described previously^83^. Briefly, cells were maintained in Dulbecco’s modified Eagle’s medium (DMEM, Invitrogen), supplemented with 15% foetal calf serum (Invitrogen), L-glutamine (Gibco) and non-essential amino acids (Gibco), betamercaptoethanol (Sigma) and leukaemia inhibitory factor (LIF; produced in-house). Experiments were performed with cells grown for several passages on plates coated with 0.2% gelatin (Sigma).

### Human cells

The human primary cell line (HMEC) was cultured in Mammary Epithelial Cell Growth Medium (Gibco MEGM) supplemented with growth factors (Gibco MEGM-SingleQuots CC-4136) BPE, hydrocortisone, hEGF, insulin and gentamicin/amphotericin-B (37°C, 7% CO_2_).

### Generation of cell lines

We tagged the endogenous *p53* and *Trim24* genes with the V5 epitope tag to facilitate detection and the FKBP12(F36V) variant in order to induce degradation upon the addition of the dTAG13 compound^33^. A V5-FKBP12(F36V) gene fragment was ordered with 100bp homology arms to the N terminus of the mouse *p53* gene, and a FKBP12(F36V)-V5 gene fragment was ordered with 100bp homology arms to the C terminus of the mouse *Trim24* gene (Twist Biosciences). A guide RNA sequence taken from the Escherichia coli genome (GTGTTGTGGACTGCGGCGGTCGG) and restriction enzyme sites were added to either end of the gene fragment via PCR. The PCR product was gel-purified using a QIAquick Gel Extraction Kit (Qiagen), digested and cloned into a donor vector. Guide RNA oligos against the N terminus of the *p53* gene (ATCCGACTGTGACTCCTCCA) and the C terminus of the *Trim24* gene (GGCGGCGTTACTTAAGCAGC) and the bacterial gRNA were ordered (Microsynth), annealed and cloned into the BsaI sites of the pC2P plasmid, which contains a Cas9-P2A-puromycin cassette. Wildtype cells (5 × 10^5) were co-transfected in suspension using Lipofectamine 3000 Reagent (Thermo Fisher Scientific) with 750 ng of the donor plasmid, 250 ng of the target gRNA plasmid and 100 ng of the bacterial gRNA plasmid. They were plated into a 6-well plate coated with 0.2% gelatin and left for 24 h. After this time the medium was replaced with puromycin (2nM) containing medium and left for another 24 h, after which the puromycin-containing medium was replaced with normal medium, and cells were left to recover for 24–48 h in normal medium. Cells were then plated for clone picking and left to grow for approximately 7 days. Individual clones were picked into a 96-well plate and genotyped by PCR. Clones that showed a homozygous knock-in at the genetic level were expanded and verified by western blot for integration of the V5 and FKBP12(F36V) before and after degradation. In addition, from the same pool of cells a subset of clones was found to have lost expression of the targeted gene due to Crispr/Cas9 cleavage without repairing-in the tagged sequence and resulting in mutations that caused a null allele of the endogenous gene. These knockout alleles were expanded and verified by western blot.

We edited the endogenously *Trim24* gene (NP 001258993.1) to harbour a CA mutation - C52A and C55A, in the RING domain of Trim24 that renders homologous Trim proteins incompatible with ubiquitin ligase activity^53^. A C52/55A gene fragment was ordered with 100bp homology arms to the RING domain of the mouse *Trim24* gene (Twist Biosciences). Gene editing was performed as for the introduction of V5 and FKBP12(F36V) sequences and using a guide RNA oligo against the RING domain of *Trim24* (ACACGGCGCAAGTGTCCAAC).

To generate addback alleles of the *Trim24* gene, RMCE was used to generate pools of cells expressing gene fragments downstream of a Cag promoter, inserted at the same genomic locus^82,84^. Geneblocks (Twist Biosciences) were ordered and cloned into a donor plasmid, including full length Trim24 (NP 001258993.1), the N terminal fragment containing the RBBC domain (AA 1-473), the N terminal deletion (AA 393-1045), the PHD null mutation variant (C807W) and variants where the PHD domain (AA793-838) was replaced with that of ING1 (AA 141-190, NP_937861.1) or TAF3 (AA 856-901, NP_114129.1). In brief, *Trim24* knockout TC-1 ES cells (background 129S6/SvEvTac) carrying an RMCE selection cassette (described previously^82^) were selected under hygromycin (250μg/ml, Roche) for 10 days. Next, 4 million cells were electroporated (Amaxa Nucleofection, Lonza) with 25μg of L1-Cag/FLAG_NLS_ORF-1L plasmid and 15μg of pIC-Cre. Negative selection with 3μM Ganciclovir (Roche) was started 2 days after transfection and continued for 10 days. Individual clones were picked into a 96-well plate and genotyped by PCR. Clones that showed an integration at the genetic level were expanded and verified by western blot. To generate GFP expressing cells, RMCE was used on the V5dTag-Trim24 mESC line to generate pools of cells expressing GFP downstream of a Cag promoter, inserted at the same genomic locus^82,84^. RMCE and clone selection were performed as described above.

### ChIP–seq

Mouse ES cells (1.5 × 10^7) were seeded into 15-cm plates the day before the experiment. Where applicable, the media was exchanged with fresh media and containing 1µM doxorubicin (#44583, Sigma-Aldrich) and/or dTAG13 compound (500nM, Tocris) in the morning four hours prior to harvesting of cells. ChIP was carried out as previously described^85^ with the following modifications: (1) chromatin was sonicated for 22 cycles of 20s on and 40s off using a Diagenode Bioruptor Pico; (2) 75μg of chromatin was used per immunoprecipitation; (3) protein A magnetic dynabeads (Thermo Fisher Scientific) were used. Immunoprecipitated DNA was subjected to library preparation (NEBNext Ultra II DNA Library Prep Kit, Illumina). In the library preparation protocol, samples were amplified using 12 PCR cycles. Libraries were sequenced on the Illumina HiSeq (50 cycles) or NextSeq (paired end 75 cycles).

### RNA-seq

Mouse ES cells were seeded into 6-well plates (2.5 × 10^5 cells/well) the day before the experiment. Where applicable, the media was exchanged with fresh media and containing 1 µM doxorubicin (#44583, Sigma-Aldrich) and/or dTAG13 compound (500nM, Tocris) for the indicated amount of time (2, 4, 8 or 12 hours) prior to harvesting. RNA was isolated using Direct-zol MicroPrep RNA Purification Kit (Zymo), following the manufacturer’s instructions. Sequencing libraries were prepared from 100 ng of purified total RNA for biological replicates using TruSeq stranded Total RNA Library Prep (Illumina). Libraries were sequenced on the Illumina HiSeq (50 cycles) or NovaSeq (paired end 100 cycles).

### ATAC-seq

ATAC-seq was performed according to the previously described protocol^83,86^ with modifications. Mouse ES cells were seeded into 6-well plates (2.5 × 10^5 cells/well) the day before the experiment and where applicable, the media was exchanged with fresh media and containing 1 µM doxorubicin (#44583, Sigma-Aldrich) to activated p53 and/or dTAG13 compound (500nM, Tocris) to degrade dTAG-tagged proteins for the indicated amount of time (1, 2, 4, 8 or 12 hours) prior to harvesting. For activation of p53 via alternative pathways, media was exchanged with fresh media containing 20 µM nutlin3a (444152, Sigma-Aldrich) for 4 hours or cells in culture media were subjected to 60Gy ionizing radiation using the automated CellRad system (Faxitron), and left for 4 hours. In brief, 5 × 10^4 cells were resuspended in 1 ml of cold ATAC-seq resuspension buffer (RSB: 10mM Tris-HCl pH 7.4, 10mM NaCl and 3mM MgCl2 in water). Cells were centrifuged at 500g for 5 minutes in a pre-chilled centrifuge (4 °C). Cell pellets were suspended in 50μl of ATAC-seq RSB containing 0.1% NP40, 0.1% Tween-20 and 0.01% digitonin by pipetting up and down three times. This cell lysis reaction was incubated on ice for 3 min. After lysis, 1 ml of ATAC-seq RSB containing 0.1% Tween-20 (without NP40 or digitonin) was added, and the tubes were inverted to mix. Nuclei were then centrifuged for 10 minutes at 500g (4 °C). Nuclei were suspended in 50μl of Tn5 transposition mix (10μl 5 × reaction buffer (50mM Tris-HCl pH 8.5, 25mM MgCl2, 50% DMF), 2.5μl transposase (100nM final), 16.5µl PBS, 0.5µl 1% digitonin, 0.5µl 10% Tween-20, 20µl water) by pipetting up and down six times. Transposition reactions were incubated at 37 °C for 30 minutes in a thermomixer with shaking at 1,000 rpm. Reactions were cleaned up using the MinElute PCR Purification Kit (Qiagen). The eluted transposed DNA was subjected to PCR amplification using Q5 High-Fidelity Polymerase (NEB) for 7 cycles. Libraries were sequenced on the Illumina NextSeq (paired end 75 cycles).

### Immunofluorescence microscopy

Mouse ES cells were seeded on poly-L-lysine coated 8-well (8*10^4 cells/well) μ-Slides (Ibidi) and left for 4 hours before the media was exchanged with fresh media and containing 1 µM doxorubicin (#44583, Sigma-Aldrich) or 20 uM nutlin3a (444152, Sigma-Aldrich) and/or dTAG13 compound (500nM, Tocris) and left for another 4 hours to activate p53 and/or degrade Trim24. Cells were fixed with 4% formaldehyde in PBS for 15 minutes, washed 3x with PBS, permeabilized in PBS with 5% foetal calf serum (Invitrogen) and 0.3% Triton X-100 for 30 minutes, then incubated overnight with antibodies (Supplementary Table 1) in PBS with 1% BSA and 0.1% Tween20. On the next day, plates were washed 3x with PBS-1% BSA and 0.1% Tween20 and incubated with secondary antibodies (1:500) in PBS-1% BSA and 0.1% Tween20 for 1 hour at room temperature. Plates were incubated for 10 minutes with PBS with Hoechst 33342 (1:500 dilution) (#134406, Thermo Fisher Scientific) and then washed 3x with PBS. Cells were imaged with a Visitron Spinning Disk W1 microscope with a 40× objective. Images were processed with ImageJ: background subtracted, nuclei segmented and mean intensity measured.

### Time lapse microscopy

Mouse ES cells were seeded on poly-L-lysine -coated 8-well (3*10^4 cells/well) μ-Slides (Ibidi) and left for 4 hours before the media was exchanged with fresh media and containing 1 µM doxorubicin (#44583, Sigma-Aldrich) or 20 uM nutlin3a (444152, Sigma-Aldrich) and/or dTAG13 compound (500nM, Tocris). Cells were kept at 37°C and 7% CO_2_ throughout the experiment, up to 40hrs post-media exchange. Cytoplasmic GFP expressing cells were imaged with a Visitron Spinning Disk W1 microscope using 488 laser and 20× objective in 10 minute intervals. At least six areas were recorded per condition. Images were processed with ImageJ. Background subtracted images were segmented separating round bright dead cells from the plate attached live cells. The mean intensities at a time point were plotted using the R package ggplot2^87^.

### Co-immunoprecipitation and Mass spectrometry

Mouse ES cells were seeded into 15cm plates (1.5 × 10^7 cells) the day before the experiment and where applicable, the media was exchanged with fresh media and containing 1 µM doxorubicin (#44583, Sigma-Aldrich) and/or dTAG13 compound (500nM, Tocris) for four hours prior to harvesting. In brief, cells were washed with PBS and harvested by treated with 0.05% trypsin. Cells were pelleted by centrifuging for 2 minutes at 300g and resuspended in 3ml PBS with 0.1% formaldehyde. After 10 minutes of crosslinking, reactions were quenched with 150µl of 2.5M glycine, vortexed briefly and left on ice for 2 minutes. Cells were pelleted by centrifuging for 3 minutes at 4°C at 500g and washed in 1.5ml PBS with 0.2% BSA, before being re-pelleted by centrifuging for 3 minutes at 4°C at 500g and resuspended in 180µl TMS buffer (10mM Tris-HCl pH 8, 1mM MgCl_2_, 1% SDS, 1× Complete Protease Inhibitor Cocktail (PIC, Roche)) with 0.4µl benzonase (E1014, Milipore) prechilled at 12°C. Samples were incubated at 12°C for 30 minutes in a thermomixer with shaking at 500 rpm, then 1570µl were added of ice-cold Dilution buffer (11.4mM Tris-HCl pH 8, 573mM NaCl, 1.14% Triton X-100, 11.46mM EDTA, 1× Complete Protease Inhibitor Cocktail (PIC, Roche)) and left on ice for 5 minutes. Samples were centrifuged at 16000g at 4°C for 5 minutes and the supernatant were transferred to new tubes with 10µl of anti-V5 mAb magnetic beads (M215-11, MBL). V5-tagged protein were bound to beads by incubating for 2 hours in overhead rotator and washed 3x with 1ml of ice-cold Wash buffer (10mM Tris-HCl pH 8, 500mM NaCl, 1% Triton X-100, 0.1% SDS, 1mM EDTA). Beads were resuspended in 5µl Digestion buffer (3M GuaHCl, 20mM EPPS pH 8.5, 10mM CAA, 5mM TCEP) + 1µl Lys-C and incubated at room temperature for 4 hours. Beads were mixed with 17uL 50mM HEPES pH 8.5, then 1µl 0.2µg/µl trypsin was added and incubated overnight at 37°C. The following morning, another 1µl of 0.2µg/µl trypsin was added and the digestion was continued for an additional 5 hours. Samples were acidified by adding 1uL of 20% TFA and sonicated in an ultrasound bath. Peptides were analysed by LC–MS/MS on an EASY-nLC 1000 (Thermo Scientific) with a two-column set-up. The peptides were applied onto a peptide μPAC™ trapping column in 0.1% formic acid, 2% acetonitrile in H2O at a constant flow rate of 5μl/min. Using a flow rate of 500 nl/min, peptides were separated at RT with a linear gradient of 3%–6% buffer B in buffer A in 4 minutes followed by a linear increase from 6 to 22% in 55 min, 22%–40% in 4 min, 40%–80% in 1 min, and the column was finally washed for 13 minutes at 80% buffer B in buffer A (buffer A: 0.1% formic acid; buffer B: 0.1% formic acid in acetonitrile) on a 50cm μPAC™ column (PharmaFluidics) mounted on an EASY-Spray™ source (Thermo Scientific) connected to an Orbitrap Fusion LUMOS (Thermo Scientific). The data were acquired using 120,000 resolution for the peptide measurements in the Orbitrap and a top T (3 s) method with HCD fragmentation for each precursor and fragment measurement in the ion trap according to the recommendation of the manufacturer (Thermo Scientific).

### Recombinant Trim24 Protein expression

Mouse full-length TRIM24 was subcloned into a pAC-derived vector^88^ containing an N-terminal Flag-tag. Recombinant protein was expressed in 8L culture of *Spodoptera frugiperda* SF9 cells using the Bac-to-Bac system (Thermo Fisher). Cells were cultured at 27°C, harvested 2 days after infection, resuspended in lysis buffer (50mM Hepes pH 8.0, 225mM NaCl, 0.1% Triton X-100, 1x protease inhibitor cocktail (Sigma)), and lysed by sonication. The supernatant was harvested and the protein was purified by FLAG-affinity purification followed by MonoQ anion exchange chromatography (GE Healthcare). Finally, TRIM24 was subjected to size exclusion chromatography (Superose 6; GE Healthcare) in SEC buffer (20mM Hepes pH 8.0, 225mM NaCl, 0.5mM TCEP, 10% glycerol). The purified protein was concentrated and stored at -80°C.

### Trim24-nuclesome binding assays

To investigate if specific histone modifications contribute to the ability of Trim24 to engage nucleosomes, we utilized the *dCypher*® approach^89^ via the Alpha® platform^90,91^. Semi-synthetic nucleosomes and controls with defined post-translational modifications were synthesized/purified/assembled using the commercial *dCypher*® and versaNuc™ services (https://www.epicypher.com/services/). Binding assays were essentially performed as previously described^89,92^ with modifications. In brief, a dCypher panel of 74 nucleosomes plus DNA/buffer controls were combined with Flag-Trim24 at 175nM (optimal screening concentration determined by initial binding measurements to candidate nucleosomes) in 10µl Binding buffer (20mM Tris pH 7.5 + 225mM NaCl + 0.01% NP-40 + 0.01% BSA + 1mM DTT) followed by a 30 minute incubation at 23°C in a 384-well plate format. A 10µl mixture of 1:400 anti-Flag antibody (Millipore Sigma F7425), 5 μg/mL Protein A Acceptor beads and 10 μg/mL Alpha Streptavidin Donor beads were added and incubated at 23°C in subdued lighting for 60 min. AlphaLISA signal was measured on a PerkinElmer 2104 EnVision (680 nm laser excitation, 570 nm emission filter ± 50 nm bandwidth). To test specific modifications (i.e. versaNucs combinatorically modified with H3K4me-0/1/2/3;H3K23-ac), binding reactions were essentially carried out and measured in the same manner, over a Flag-Trim24 dilution series of 0.4µM to 0.39nM. All measurements were performed in duplicate and mean signal values are shown with error bars indicating standard error. Relative EC50 (EC50rel) values were computed using a four-parameter logistical (4PL) model in GraphPad Prism 8 as previously described^93,94^.

### Computational analyses ChIPseq

ChIPseq datasets were aligned to either the mm10 mouse or hg19 human assembly using the QuasR^95^ Bioconductor package that utilizes Bowtie^96^ (RBowtie package). Alignments were performed with the default settings and allowing for uniquely mapping reads. Peak calling on all datasets were performed with MACS2^97^ (version 2.1.3.3) using the callpeak argument with default settings and specifying the genome size with -g mm or -hs for mouse or human, respectively. Peaks were called for mouse ChIPseq datasets using matched IgG ChIPseq datasets as controls. Peaks were called for human ChIPseq datasets using either matched IgG ChIPseq, or matched chromatin input sequencing datasets as controls. Peaks from replicate datasets and across different samples were unified by sorting and merging overlapping regions using the bedtools^98^ (version v2.25.0) ‘sort’ and ‘merge’ functions with default settings, as in the case of human p53 ChIPseq (Supplementary Fig. 2) and Trim24-addback ChIPseq (Supplementary Fig. 10b-c). Motif enrichments on individual ChIPseq datasets were performed using the HOMER^99^ software, whereby each peak set were ranked according to MACS2-definned *P* values and the top 500 peaks were used. The HOMER findMotifsGenome.pl function was run on these top peaks, using the -len argument to search for motifs of 10, 12, 14, 16 bp in length and the -size argument to search for motifs within 250bp around the peak centre. Both known and *de novo* motif finding was performed and are reported as indicated within figures and visualized by using HOMER nucleotide frequencies and the Bioconductor SeqLogo^100^ R package. Read counts were generated over defined genomic regions (i.e. peak regions) using the QuasR^95^ function qCount after removing blacklisted regions^101^, with default parameters and shifting the reads 80bp, which was approximately half the size of ChIPseq library fragments. For datasets with paired end sequencing, this was instead shifted to half of the fragment length. Briefly, counts were normalized between datasets being compared, a pseudo-count of 8 was added and data were log2-transformed. Normalization was performed by multiplying counts by a scaling factor that was determined by the library with the lowest number of mapped reads between the datasets, i.e., scaled down to the smaller library: Scaling factor χ= min(Sample1, Sample2… Sample χ)/Sample χ, where Sample 1, Sample2, Sample χ are the total number of mapped reads in each respective sample. The pseudo-count of 8 was used to decrease noise at low read counts between samples. Enrichments of log2 ChIPseq readcounts were calculated by subtracting the matched log2 counts from corresponding control datasets. Similarly, changes in binding were calculated by the difference in log2 ChIPseq readcounts between datasets. To define a reference set of p53 and Trim24 peaks from the replicate experiments in parental mESC line with or without p53 activation, the consensus peak calling was carried out as follows: MACS2 peaks from individual replicates were assembled using R package DiffBind^102^ (v3.2.4) to generate a non-overlapping set of genomic peaks. As described in the Encode project^103^, Irreproducibility Discovery Rate^104^ (IDR) analysis was applied to define a reliable set of consensus peaks using a threshold of IDR <0.01, average ChIPseq enrichment > 2 and a minimum ChIPseq enrichment of 0.5 in each replicate. Heatmaps were generated by counting the 5’ positions of mapped reads relative to defined genomic regions (i.e. peak regions) using the QuasR^95^ function qProfile and visualized using the EnrichedHeatmap^105^ Bioconductor package. In brief, qProfile was run with default parameters, for 1kb regions centred on the middle of peak regions and shifting the reads 80bp, which was approximately half the size of ChIPseq library fragments, or to the fragment midpoint for paired-end libraries. Resulting counts/peak region were scaled by 1e6/total reads in each sample and multiplied by 1e3, then smoothed by calculating a running mean of 20bp across the normalized counts. These were converted into normalized matrixes, replicates averaged, and visualized using the as.normalizedMatrix and EnrichedHeatmap functions from the EnrichedHeatmap^105^ R package. Colour scales were implemented manually based on enrichment values and using the colorRamp2 function within the Circlize^106^ R package. For histone mark metaprofiles, profiles from each mark were normalized by dividing their maximum value or by 1.5 if the maximum was < 1.5^107^. To cluster p53 peaks by enrichments for histone marks, k-means clustering on a histone datasets were performed using the kmeans function from the R stats^108^ package with the arguments centres=4 and nstart = 10. Publicly available datasets utilized in this study are as indicated (Supplementary Table 2).

### RNAseq

RNAseq reads were mapped to mm10 using STAR^109^ aligner (version 2.5.2b). Alignments were performed with otherwise default settings and using the arguments –outFilterType BySJout, -- outFilterMultimapNmax 20, --alignSJoverhangMin 8, --alignSJDBoverhangMin 3, --alignIntronMin 20, -- outSAMmultNmax 1, allowing for up to 20 matches for a multimapping read and outputting one at random in these cases. Resulting alignment files were indexed using Samtools^110^ (version 1.2) using default parameters. Alignments overlapping protein coding genes were counted using the Rsubread^111^ Bioconductor package. In brief, the featureCounts function from Rsubread was run with otherwise default settings and with the gene annotation from the M25 GENCODE^112^ release, and with GTF.attrType = “gene_name” to group gene features (e.g. exons). GENCODE annotations (i.e., protein_coding) were used to retain only those resulting counts for protein coding genes and excluding predicted genes (i.e. beginning ‘Gm’ id number). Count matrixes were generated and TMM-normalized by the DGEList and calcNormFactors functions from the edgeR^113^ package using default settings. Finally, counts per million (CPM) were generated using the cpm function from edgeR and with prior.count = 8 and log = True arguments, to add a pseudocount of 8 and to log2 transform the data. Analysis to determine statistical significance in differential expression between groups of samples were performed using edgeR. Briefly, read counts over protein coding genes were generated and normalized as above for specific samples and a design matrix was generated for these in the groups being compared using the model.matrix function. Significance and foldchange estimates were generated by the estimateDisp, glmQLFit and glmQLFTest and predFC functions in the edgeR package followed by multiple testing correction with Benjamini-Hochberg method. Genes were considered differentially expressed with an FDR<0.01 and an absolute log2 foldchange of at least 0.75. A consensus set of Trim24-target genes in the Trim24 degron line were nominated as those that were differentially expressed upon degradation of Trim24 at the 4-hour timepoint of p53 activation; a later time point (8 hours) was well correlated, and effects were largely unidirectional, i.e., increased expression compared to Trim24-expressing cells. Genes increasing in expression at this point were enriched in Trim24 and p53 binding in promoters (*P* value < 2.2e-16; Odds ratio 5.4) and at the 12-hour point, expression of these were less well-correlated and a roughly even number of genes further increased and decreased, likely due to secondary effects on gene expression as a result of miss-expression of the primary targets (Supplementary Fig. 12b). This consensus set therefore represents a conservative estimate of direct Trim24-target genes. To test the distance relationships between differentially expressed genes and genomic regions (i.e., p53 binding peak regions), the annotatePeak function from the ChIPseeker^114^ package was used to identify genomic regions within or nearest gene. Briefly, annotatePeak was run with default arguments and utilizing the TxDb.Mmusculus.UCSC.mm10.knownGene^115^ and org.Mm.eg.db^116^ annotation packages. Functional enrichment analysis of Trim24-target genes was carried out by the R package ‘clusterProfiler’ using Gene Ontology (GO) and KEGG annotations. The featureCounts tool^117^ (v2.0.0) was employed to determine RNAseq counts over repetitive elements (e.g. Lines, Sines, LTR-containing retrotransposons etc.) using RepeatMasker^118^ annotations (version 4.1.2) in Gtf format obtained from UCSC^119^ genome browser database (mm10). edgeR, as described above, was used to assess the differential expression of repetitive elements upon Trim24 degradation. To identify p53 motifs in repeat LTR elements, the Repbase^120,121^ (version 20.02) consensus sequences were scanned for the p53 weight matrix derived from the Jaspar MA0106.3 p53 motif using the matchPWM function from the Biostrings^122^ R package. Matching sequences were determined by requiring a log2-odds score of at least 10 over a uniform background.

### ATACseq

ATACseq reads were trimmed using cutadapt^123^ (version 2.5), with parameters -a CTGTCTCTTATACACA -A CTGTCTCTTATACACA -m 10 --overlap = 1 and then mapped to mm10 using QuasR^95^ with defaults settings. To determine read counts over genomic regions (e.g., ChIPseq peaks), reads were first counted over all regions using the QuasR^95^ function qCount with default parameters. Counts between samples were normalized using edgeR. In brief, ATACseq peaks for samples being compared were first generated by MACS2 as described above for ChIPseq and any peaks overlapping in at least two samples were retained and merged using the reduce function of the GenomicRanges^124^ R package, excluding those overlapping mm10 blacklisted regions^101^. Reads counts at merged peaks were then determined by qCount with default parameters. The resulting count data were then merged with counts over peak regions (i.e. ChIPseq peaks), and the combined set were TMM-normalized by the DGEList and calcNormFactors functions from the edgeR^113^ package using default settings. Finally, counts per million (CPM) were generated using the cpm function from edgeR and with prior.count = 8 and log = True arguments, to add a pseudocount of 8 and to log2 transform the data. This ensures that counts in regions of interest (i.e., ChIPseq peaks) were normalized against all accessible chromatin sites, the majority of which are not expected to change between samples. ATACseq counts in ATACseq peaks were generated in the same manner. Analysis to determine statistical significance in differential accessibility between groups of samples were performed using edgeR. Briefly, read counts over regions (i.e., ATACseq peaks) were generated and normalized as above for specific samples and a design matrix was generated for these in the groups being compared using the model.matrix function. Significance and foldchange estimates were generated by the estimateDisp, glmQLFit and glmQLFTest and predFC functions in the edgeR package. Regions were considered differentially accessible with an FDR < 0.05 and foldchange > 0.5. The monaLisa^125^ R package was used to identify motifs that were enriched in regions that change between samples. Briefly, foldchanges between samples were generated from subtracting the replicate-averaged counts in genomic regions (i.e. ATACseq peaks) and genomic sequence within peaks resized to the median with of all peaks being considered around the peak centre, using the trim and getSeq tools from the GenomicRanges^124^ and Biostrings^122^ R libraries, respectively. Fold changes between regions were binned using the bin function in the monaLisa library with default parameters and minAbsX = 1 to set the minimal absolute value for log2 changes, and the number of regions per changing bin set using the nElements argument and as described in each figure. Enrichment of motifs was performed with the calcBinnedMotifEnrR function, using motifs from the JASPAR2020^126^ database, motifs were included if they were at least -log10 adjusted *P* value >4 and log2 enriched > 0.5 within bins excluding the non-changing centre bin. For visualization, enriched motifs were clustered by similarity using the motifSimilarity and hclust tools from the monaLisa^125^ and stats^108^ packages. Heatmaps were generated by counting the midpoint positions of mapped fragments relative to defined genomic regions (i.e. peak regions) using the QuasR^95^ function qProfile and visualized using the EnrichedHeatmap^105^ Bioconductor package. In brief, qProfile was run with default parameters and with the parameters shift=“halfInsert” and useRead=“first”, for 1kb regions centered on the middle of peak regions. Resulting counts/peak region were scaled by 1e6/total reads in each sample and multiplied by 1e3, then smoothed by calculating a running mean of 20bp across the normalized counts. These were converted into normalized matrixes, replicates averaged, and visualized using the as.normalizedMatrix and EnrichedHeatmap functions from the EnrichedHeatmap^105^ R package. Colour scales were implemented manually based on enrichment values and using the colorRamp2 function within the Circlize^106^ R package.

### DNaseSeq

DNaseSeq datasets were aligned to the mm10 mouse assembly using the QuasR^95^ Bioconductor package that utilizes Bowtie^96^ (RBowtie package). Alignments were performed with the default settings and allowing for uniquely mapping reads. Read counts over defined genomic regions (i.e., peak regions) were performed using the QuasR^95^ function qCount with default parameters and shifting the reads 20bp. Briefly, counts were normalized between datasets being compared, a pseudo-count of 8 was added and data were log2-transformed. Normalization was performed by multiplying counts by a scaling factor that was determined by the library with the lowest number of mapped reads between the datasets, i.e., scaled down to the smaller library: Scaling factor χ= min(Sample1, Sample2… Sample χ)/Sample χ, where Sample 1, Sample2, Sample χ are the total number of mapped reads in each respective sample. The pseudo-count of 8 was used to decrease noise at low read counts between samples. The mean counts between replicates at genomic regions (i.e. peaks) were considered and a cut-off of > log2 6 counts was used to define open and closed regions, as this reflected an inflection point in the data using qqnorm and qqline functions from the stats^108^ R package. Heatmaps were generated by counting the 5’ positions of mapped reads relative to defined genomic regions (i.e. peak regions) using the QuasR^95^ function qProfile and visualized using the EnrichedHeatmap^105^ Bioconductor package. In brief, qProfile was run with default parameters, for 1kb regions centered on the middle of peak regions. Resulting counts/peak region were scaled by 1e6/total reads in each sample and multiplied by 1e3, then smoothed by calculating a running mean of 20bp across the normalized counts. These were converted into normalized matrixes, replicates averaged, and visualized using the as.normalizedMatrix and EnrichedHeatmap functions from the EnrichedHeatmap^105^ R package. Colour scales were implemented manually based on enrichment values and using the colorRamp2 function within the Circlize^106^ R package. Publicly available human Encode datasets (i.e. mapped bam files) utilized in this study are as indicated (Supplementary Table 2).

The quality control of sequencing data was carried out by Qualimap^127^ (v.2.2.1) using ‘bamqc’ and ‘rnaseq’ modes. The quality of ChIPseq, ATACseq and RNAseq datasets were further assessed by R ChIPQC^128^ package (v1.28.0) (Supplementary Table S3).

### IP-MS protein enrichment analysis

Protein identification and relative quantification was performed with MaxQuant v.1.5.3.8 using Andromeda as the search engine^129^ and label-free quantification (LFQ)^130^. The mouse subset of the UniProt v.2019_04 combined with the contaminant database from MaxQuant was searched and the protein and peptide FDR were set to 1% and 0.1% FDR. The combined intensity of peptides of proteins were imported into R and values were normalized between samples by dividing values of each by the sum of all values within a sample, then multiplying these by the sum of all values of the sample with the lowest sum values. Thus, values were scaled down to the sample with the least signal. Data were log2 transformed after dividing samples by 2^20 and adding a pseudocount of 5 in order to stabilize variance of the data. p53 enriched samples were compared to datasets generated by IP-MS in the p53 degron line after degradation (i.e., mock IP) and significance estimates were generated using the eBayes function in the limma^131^ R package. Briefly, the lmFit, makeContrasts and contrasts.fit tools from limma were used with default parameters to generate linear models and estimate coefficients and standard errors from samples grouped by treatment condition (i.e., p53 degraded, untreated, p53-activated). Finally, adjusted *P* values were generated by the eBayes and topTable functions from limma and proteins with < 0.01 adjusted *P* value were considered significantly enriched. As Fkbp1a peptides constitute the degron tag that was added to the endogenous p53 gene in this experiment, the Fkbp1a protein was manually removed from the list of enriched proteins.

### Estimate of coefficient contributions to linear regression models

To estimate coefficient contributions to Trim24 addback binding datasets, the lm tool from the stats^108^ R package was used to generate linear regression models and these were visualized using the plot_summs function within the jtools^132^ R package. Briefly, lm was run using replicate-averaged enrichments for Trim24-addback variant ChIPseq datasets, using default parameters and with p53 enrichment and enrichment of histone marks as independent variables. The plot_summs function was run with default parameters on the resulting fits using default parameters in order to visualize estimates of coefficient contributions to models.

### Correlation heatmaps

Correlations heatmaps to group effects and show reproducibility were generated with the cor and aHeatmap functions from the stats^108^ and NMF^133^ packages. Where applicable in the figures, dataset counts, either averages across samples or individual replicates, were used within the cor function to generate Pearson correlation coefficients and these were used directly within aHeatmap that computes a dendrogram from hierarchical clustering.

In all box plots, middle points correspond to median, boxes to first and third quartile and whiskers to 1.5 multiplied by the interquartile range. Notches, where indicated, extend to ± 1.58 × (interquartile range/square_root(n)). Whiskers correspond to the maximum and minimum distribution values after removal of outliers, where outliers were defined as more than 1.5 × (interquartile range) away from the box. Pearson correlation coefficients were calculated using the R function cor with default parameters.

For ChIPseq, Illumina RTA 1.18.64 and bcl2fastq2 v2.17 was used for basecalling and demultiplexing for single-read experiments, Illumina RTA 2.4.11 and bcl2fastq2 v2.17 was used for basecalling and demultiplexing for paired-read experiments. For RNAseq, Illumina RTA 1.18.64 and bcl2fastq2 v2.17 was used for basecalling and demultiplexing samples generated by Illumina HiSeq sequencing. Illumina RTA 3.4.4 and bcl2fastq2 v2.20 was used for basecalling and demultiplexing samples generated by Illumina NovaSeq sequencing. For ATACseq, Illumina RTA 2.4.11 and bcl2fastq2 v2.17 was used for basecalling and demultiplexing.

## References

1. Jenuwein, T. & Allis, C.D. Translating the histone code. Science 293, 1074–80 (2001).

2. Morgan, M.A.J. & Shilatifard, A. Reevaluating the roles of histone-modifying enzymes and their associated chromatin modifications in transcriptional regulation. Nature Genetics 52, 1271–1281 (2020).

3. Kouzarides, T. Chromatin modifications and their function. Cell 128, 693–705 (2007).

4. Zhang, T., Cooper, S. & Brockdorff, N. The interplay of histone modifications – writers that read. EMBO reports 16, 1467–1481 (2015).

5. Soto, L.F. et al. Compendium of human transcription factor effector domains. Molecular Cell 82, 514–526 (2022).

6. Matsumoto, S. et al. DNA damage detection in nucleosomes involves DNA register shifting. Nature 571, 79–84 (2019).

7. Michael, A.K. & Thomä, N.H. Reading the chromatinized genome. Cell 184, 3599–3611 (2021).

8. Henikoff, S. & Shilatifard, A. Histone modification: cause or cog? Trends in Genetics 27, 389–396 (2011).

9. Simon, M. et al. Histone fold modifications control nucleosome unwrapping and disassembly. Proc Natl Acad Sci U S A 108, 12711–6 (2011).

10. Dann, G.P. et al. ISWI chromatin remodellers sense nucleosome modifications to determine substrate preference. Nature 548, 607–611 (2017).

11. Mashtalir, N. et al. Chromatin landscape signals differentially dictate the activities of mSWI/SNF family complexes. Science 373, 306–315 (2021).

12. Thurman, R.E. et al. The accessible chromatin landscape of the human genome. Nature 489, 75–82 (2012).

13. Kundaje, A. et al. Integrative analysis of 111 reference human epigenomes. Nature 518, 317–330 (2015).

14. Zhou, V.W., Goren, A. & Bernstein, B.E. Charting histone modifications and the functional organization of mammalian genomes. Nat Rev Genet 12, 7–18 (2011).

15. Barisic, D., Stadler, M.B., Iurlaro, M. & Schübeler, D. Mammalian ISWI and SWI/SNF selectively mediate binding of distinct transcription factors. Nature 569, 136–140 (2019).

16. Srivastava, D. & Mahony, S. Sequence and chromatin determinants of transcription factor binding and the establishment of cell type-specific binding patterns. Biochimica et Biophysica Acta (BBA) Gene Regulatory Mechanisms 1863, 194443 (2020).

17. Zaret, K.S. & Carroll, J.S. Pioneer transcription factors: establishing competence for gene expression. Genes Dev 25, 2227–41 (2011).

18. Dodonova, S.O., Zhu, F., Dienemann, C., Taipale, J. & Cramer, P. Nucleosome-bound SOX2 and SOX11 structures elucidate pioneer factor function. Nature 580, 669–672 (2020).

19. Michael, A.K. et al. Mechanisms of OCT4-SOX2 motif readout on nucleosomes. Science 368, 1460–1465 (2020).

20. Sullivan, K.D., Galbraith, M.D., Andrysik, Z. & Espinosa, J.M. Mechanisms of transcriptional regulation by p53. Cell Death & Differentiation 25, 133–143 (2018).

21. Yan, H. et al. p53 is active in murine stem cells and alters the transcriptome in a manner that is reminiscent of mutant p53. Cell Death & Disease 6, e1662–e1662 (2015).

22. Peng, T. et al. STARR-seq identifies active, chromatin-masked, and dormant enhancers in pluripotent mouse embryonic stem cells. Genome Biology 21, 243 (2020).

23. Younger, S.T. & Rinn, J.L. p53 regulates enhancer accessibility and activity in response to DNA damage. Nucleic Acids Research 45, 9889–9900 (2017).

24. Sammons, M.A., Zhu, J., Drake, A.M. & Berger, S.L. TP53 engagement with the genome occurs in distinct local chromatin environments via pioneer factor activity. Genome research 25, 179–188 (2015).

25. Kribelbauer, J.F. et al. Quantitative Analysis of the DNA Methylation Sensitivity of Transcription Factor Complexes. Cell Rep 19, 2383–2395 (2017).

26. Stadler, M.B. et al. DNA-binding factors shape the mouse methylome at distal regulatory regions. Nature 480, 490–5 (2011).

27. Buenrostro, J.D., Wu, B., Chang, H.Y. & Greenleaf, W.J. ATAC-seq: A Method for Assaying Chromatin Accessibility Genome-Wide. Curr Protoc Mol Biol 109, 21.29.1-21.29.9 (2015).

28. Song, L. & Crawford, G.E. DNase-seq: a high-resolution technique for mapping active gene regulatory elements across the genome from mammalian cells. Cold Spring Harb Protoc 2010, pdb.prot5384 (2010).

29. Tacar, O., Sriamornsak, P. & Dass, C.R. Doxorubicin: an update on anticancer molecular action, toxicity and novel drug delivery systems. Journal of Pharmacy and Pharmacology 65, 157–170 (2012).

30. Lee, K.-H. et al. A genomewide study identifies the Wnt signaling pathway as a major target of p53 in murine embryonic stem cells. Proceedings of the National Academy of Sciences 107, 69–74 (2010).

31. Domcke, S. et al. Competition between DNA methylation and transcription factors determines binding of NRF1. Nature 528, 575–9 (2015).

32. Wang, B., Xiao, Z., Ko, H.L. & Ren, E.C. The p53 response element and transcriptional repression. Cell Cycle 9, 870–9 (2010).

33. Nabet, B. et al. The dTAG system for immediate and target-specific protein degradation. Nature chemical biology 14, 431–441 (2018).

34. Grand, R.S. et al. BANP opens chromatin and activates CpG-island-regulated genes. Nature 596, 133–137 (2021).

35. Allton, K. et al. Trim24 targets endogenous p53 for degradation. Proceedings of the National Academy of Sciences 106, 11612–11616 (2009).

36. Tsai, W.-W. et al. TRIM24 links a non-canonical histone signature to breast cancer. Nature 468, 927–932 (2010).

37. Vassilev, L.T. et al. In Vivo Activation of the p53 Pathway by Small-Molecule Antagonists of MDM2. Science 303, 844–848 (2004).

38. Lowe, S.W., Schmitt, E.M., Smith, S.W., Osborne, B.A. & Jacks, T. p53 is required for radiation-induced apoptosis in mouse thymocytes. Nature 362, 847–9 (1993).

39. Champagne, K.S. & Kutateladze, T.G. Structural insight into histone recognition by the ING PHD fingers. Current drug targets 10, 432–441 (2009).

40. Vermeulen, M. et al. Selective anchoring of TFIID to nucleosomes by trimethylation of histone H3 lysine 4. Cell 131, 58–69 (2007).

41. Peña, P.V. et al. Histone H3K4me3 Binding Is Required for the DNA Repair and Apoptotic Activities of ING1 Tumor Suppressor. Journal of Molecular Biology 380, 303–312 (2008).

42. Padeken, J., Methot, S.P. & Gasser, S.M. Establishment of H3K9-methylated heterochromatin and its functions in tissue differentiation and maintenance. Nature Reviews Molecular Cell Biology (2022).

43. Blackledge, N.P. & Klose, R.J. The molecular principles of gene regulation by Polycomb repressive complexes. Nature Reviews Molecular Cell Biology 22, 815–833 (2021).

44. Herquel, B. et al. Trim24-repressed VL30 retrotransposons regulate gene expression by producing noncoding RNA. Nat Struct Mol Biol 20, 339–46 (2013).

45. Wylie, A. et al. p53 genes function to restrain mobile elements. Genes Dev 30, 64–77 (2016).

46. Chang, N.T., Yang, W.K., Huang, H.C., Yeh, K.W. & Wu, C.W. The transcriptional activity of HERV-I LTR is negatively regulated by its cis-elements and wild type p53 tumor suppressor protein. J Biomed Sci 14, 211–22 (2007).

47. Leonova, K.I. et al. p53 cooperates with DNA methylation and a suicidal interferon response to maintain epigenetic silencing of repeats and noncoding RNAs. Proceedings of the National Academy of Sciences 110, E89–E98 (2013).

48. Clària, J. Cyclooxygenase-2 biology. Curr Pharm Des 9, 2177–90 (2003).

49. Corcoran, C.A., He, Q., Huang, Y. & Sheikh, M.S. Cyclooxygenase-2 interacts with p53 and interferes with p53-dependent transcription and apoptosis. Oncogene 24, 1634–1640 (2005).

50. Li, Y. et al. Structural insights into the TRIM family of ubiquitin E3 ligases. Cell Res 24, 762–5 (2014).

51. Randolph, K., Hyder, U. & D’Orso, I. KAP1/TRIM28: Transcriptional Activator and/or Repressor of Viral and Cellular Programs? Front Cell Infect Microbiol 12, 834636 (2022).

52. Rowe, H.M. et al. KAP1 controls endogenous retroviruses in embryonic stem cells. Nature 463, 237–40 (2010).

53. Dupont, S. et al. Germ-layer specification and control of cell growth by Ectodermin, a Smad4 ubiquitin ligase. Cell 121, 87–99 (2005).

54. Shen, Y. et al. A map of the cis-regulatory sequences in the mouse genome. Nature 488, 116–120 (2012).

55. Groner, A.C. et al. TRIM24 Is an Oncogenic Transcriptional Activator in Prostate Cancer. Cancer Cell 29, 846–858 (2016).

56. Herquel, B. et al. Transcription cofactors TRIM24, TRIM28, and TRIM33 associate to form regulatory complexes that suppress murine hepatocellular carcinoma. Proceedings of the National Academy of Sciences of the United States of America 108, 8212–8217 (2011).

57. Li, H. et al. Overexpression of TRIM24 correlates with tumor progression in non-small cell lung cancer. PLoS One 7, e37657 (2012).

58. Zhang, L.H. et al. TRIM24 promotes glioma progression and enhances chemoresistance through activation of the PI3K/Akt signaling pathway. Oncogene 34, 600–10 (2015).

59. Shah, V.V. et al. Mammary-specific expression of Trim24 establishes a mouse model of human metaplastic breast cancer. Nature Communications 12, 5389 (2021).

60. Lv, D. et al. TRIM24 is an oncogenic transcriptional co-activator of STAT3 in glioblastoma. Nature Communications 8, 1454 (2017).

61. Appikonda, S., Thakkar, K.N. & Barton, M.C. Regulation of gene expression in human cancers by TRIM24. Drug Discovery Today: Technologies 19, 57–63 (2016).

62. Gechijian, L.N. et al. Functional TRIM24 degrader via conjugation of ineffectual bromodomain and VHL ligands. Nature Chemical Biology 14, 405–412 (2018).

63. Alexanian, A. & Sorokin, A. Cyclooxygenase 2: protein-protein interactions and posttranslational modifications. Physiol Genomics 49, 667–681 (2017).

64. FitzGerald, G.A. & Patrono, C. The coxibs, selective inhibitors of cyclooxygenase-2. N Engl J Med 345, 433–42 (2001).

65. Williams, C.S., Mann, M. & DuBois, R.N. The role of cyclooxygenases in inflammation, cancer, and development. Oncogene 18, 7908–16 (1999).

66. Marnett, L.J. The COXIB experience: a look in the rearview mirror. Annu Rev Pharmacol Toxicol 49, 265–90 (2009).

67. Hashemi Goradel, N., Najafi, M., Salehi, E., Farhood, B. & Mortezaee, K. Cyclooxygenase-2 in cancer: A review. J Cell Physiol 234, 5683–5699 (2019).

68. Khetchoumian, K. et al. Loss of Trim24 (Tif1alpha) gene function confers oncogenic activity to retinoic acid receptor alpha. Nat Genet 39, 1500–6 (2007).

69. Kikuchi, M. et al. TRIM24 mediates ligand-dependent activation of androgen receptor and is repressed by a bromodomain-containing protein, BRD7, in prostate cancer cells. Biochim Biophys Acta 1793, 1828–36 (2009).

70. Matsui, T. et al. Proviral silencing in embryonic stem cells requires the histone methyltransferase ESET. Nature 464, 927–931 (2010).

71. Isbel, L. et al. Trim33 Binds and Silences a Class of Young Endogenous Retroviruses in the Mouse Testis; a Novel Component of the Arms Race between Retrotransposons and the Host Genome. PLoS Genet 11, e1005693 (2015).

72. Soufi, A., Donahue, G. & Zaret, K.S. Facilitators and impediments of the pluripotency reprogramming factors’ initial engagement with the genome. Cell 151, 994–1004 (2012).

73. Chen, D. et al. A Multidimensional Characterization of E3 Ubiquitin Ligase and Substrate Interaction Network. iScience 16, 177–191 (2019).

74. O’Connor, H.F. & Huibregtse, J.M. Enzyme-substrate relationships in the ubiquitin system: approaches for identifying substrates of ubiquitin ligases. Cell Mol Life Sci 74, 3363–3375 (2017).

75. Zhu, Q. et al. TRIM24 facilitates antiviral immunity through mediating K63-linked TRAF3 ubiquitination. J Exp Med 217(2020).

76. Komander, D. & Rape, M. The Ubiquitin Code. Annual Review of Biochemistry 81, 203–229 (2012).

77. Lill, N.L., Grossman, S.R., Ginsberg, D., DeCaprio, J. & Livingston, D.M. Binding and modulation of p53 by p300/CBP coactivators. Nature 387, 823–7 (1997).

78. Barlev, N.A. et al. Acetylation of p53 activates transcription through recruitment of coactivators/histone acetyltransferases. Mol Cell 8, 1243–54 (2001).

79. Ito, M. et al. Identity between TRAP and SMCC Complexes Indicates Novel Pathways for the Function of Nuclear Receptors and Diverse Mammalian Activators. Molecular Cell 3, 361–370 (1999).

80. An, W., Kim, J. & Roeder, R.G. Ordered Cooperative Functions of PRMT1, p300, and CARM1 in Transcriptional Activation by p53. Cell 117, 735–748 (2004).

81. Gamper Armin, M. & Roeder Robert, G. Multivalent Binding of p53 to the STAGA Complex Mediates Coactivator Recruitment after UV Damage. Molecular and Cellular Biology 28, 2517–2527 (2008).

82. Lienert, F. et al. Identification of genetic elements that autonomously determine DNA methylation states. Nat Genet 43, 1091–7 (2011).

83. Mohn, F. et al. Lineage-specific polycomb targets and de novo DNA methylation define restriction and potential of neuronal progenitors. Mol Cell 30, 755–66 (2008).

84. Feng, Y.Q. et al. Site-specific chromosomal integration in mammalian cells: highly efficient CRE recombinase-mediated cassette exchange. J Mol Biol 292, 779–85 (1999).

85. Weber, M. et al. Distribution, silencing potential and evolutionary impact of promoter DNA methylation in the human genome. Nat Genet 39, 457–66 (2007).

86. Corces, M.R. et al. An improved ATAC-seq protocol reduces background and enables interrogation of frozen tissues. Nature Methods 14, 959–962 (2017).

87. Villanueva, R.A.M. & Chen, Z.J. ggplot2: elegant graphics for data analysis. (Taylor & Francis, 2019).

88. Abdulrahman, W. et al. A set of baculovirus transfer vectors for screening of affinity tags and parallel expression strategies. Anal Biochem 385, 383–5 (2009).

89. Marunde, M.R., Popova, I.K., Weinzapfel, E.N. & Keogh, M.C. The dCypher Approach to Interrogate Chromatin Reader Activity Against Posttranslational Modification-Defined Histone Peptides and Nucleosomes. Methods Mol Biol 2458, 231–255 (2022).

90. Eglen, R.M. et al. The use of AlphaScreen technology in HTS: current status. Current chemical genomics 1, 2–10 (2008).

91. Quinn, A.M. et al. A homogeneous method for investigation of methylation-dependent protein– protein interactions in epigenetics. Nucleic Acids Research 38, e11–e11 (2009).

92. Jain, K. et al. Characterization of the plant homeodomain (PHD) reader family for their histone tail interactions. Epigenetics & chromatin 13, 3–3 (2020).

93. Morgan, M.A.J. et al. A trivalent nucleosome interaction by PHIP/BRWD2 is disrupted in neurodevelopmental disorders and cancer. Genes Dev 35, 1642–1656 (2021).

94. Marunde, M.R. et al. Nucleosome conformation dictates the histone code. bioRxiv, 2022.02.21.481373 (2022).

95. Gaidatzis, D., Lerch, A., Hahne, F. & Stadler, M.B. QuasR: quantification and annotation of short reads in R. Bioinformatics 31, 1130–1132 (2015).

96. Langmead, B., Trapnell, C., Pop, M. & Salzberg, S.L. Ultrafast and memory-efficient alignment of short DNA sequences to the human genome. Genome Biology 10, R25 (2009).

97. Zhang, Y. et al. Model-based Analysis of ChIP-Seq (MACS). Genome Biology 9, R137 (2008).

98. Quinlan, A.R. & Hall, I.M. BEDTools: a flexible suite of utilities for comparing genomic features. Bioinformatics 26, 841–2 (2010).

99. Heinz, S. et al. Simple Combinations of Lineage-Determining Transcription Factors Prime cis-Regulatory Elements Required for Macrophage and B Cell Identities. Molecular Cell 38, 576–589 (2010).

100. Bembom, O. & Ivanek, R. seqLogo: Sequence logos for DNA sequence alignments. (2021).

101. Amemiya, H.M., Kundaje, A. & Boyle, A.P. The ENCODE Blacklist: Identification of Problematic Regions of the Genome. Scientific Reports 9, 9354 (2019).

102. Ross-Innes, C.S. et al. Differential oestrogen receptor binding is associated with clinical outcome in breast cancer. Nature 481, 389–393 (2012).

103. Kundaje, A. et al. ENCODE: TF ChIP-seq peak calling using the Irreproducibility Discovery Rate (IDR) framework. (2014).

104. Li, Q., Brown, J.B., Huang, H. & Bickel, P.J. Measuring reproducibility of high-throughput experiments. The Annals of Applied Statistics 5, 1752-1779, 28 (2011).

105. Gu, Z., Eils, R., Schlesner, M. & Ishaque, N. EnrichedHeatmap: an R/Bioconductor package for comprehensive visualization of genomic signal associations. BMC Genomics 19, 234 (2018).

106. Gu, Z., Gu, L., Eils, R., Schlesner, M. & Brors, B. circlize implements and enhances circular visualization in R. Bioinformatics 30, 2811–2812 (2014).

107. Iurlaro, M. et al. Mammalian SWI/SNF continuously restores local accessibility to chromatin. Nature Genetics 53, 279–287 (2021).

108. Team, R.C. R: A Language and Environment for Statistical Computing. (2022).

109. Dobin, A. et al. STAR: ultrafast universal RNA-seq aligner. Bioinformatics 29, 15–21 (2013).

110. Li, H. et al. The Sequence Alignment/Map format and SAMtools. Bioinformatics 25, 2078–2079 (2009).

111. Liao, Y., Smyth, G.K. & Shi, W. The R package Rsubread is easier, faster, cheaper and better for alignment and quantification of RNA sequencing reads. Nucleic Acids Research 47, e47–e47 (2019).

112. Frankish, A. et al. GENCODE 2021. Nucleic Acids Research 49, D916–D923 (2021).

113. Robinson, M.D., McCarthy, D.J. & Smyth, G.K. edgeR: a Bioconductor package for differential expression analysis of digital gene expression data. Bioinformatics (Oxford, England) 26, 139–140 (2010).

114. Yu, G., Wang, L.-G. & He, Q.-Y. ChIPseeker: an R/Bioconductor package for ChIP peak annotation, comparison and visualization. Bioinformatics 31, 2382–2383 (2015).

115. Maintainer, B.C.T.a.B.P. TxDb.Mmusculus.UCSC.mm10.knownGene: Annotation package for TxDb object(s). (2019).

116. Carlson, M. org.Mm.eg.db: Genome wide annotation for Mouse. (2021).

117. Liao, Y., Smyth, G.K. & Shi, W. featureCounts: an efficient general purpose program for assigning sequence reads to genomic features. Bioinformatics 30, 923–30 (2014).

118. Smit, A., Hubley, R. & Green, P. RepeatMasker Open-4.0. (2013-2015).

119. Navarro Gonzalez, J. et al. The UCSC Genome Browser database: 2021 update. Nucleic Acids Res 49, D1046–d1057 (2021).

120. Jurka, J., Walichiewicz, J. & Milosavljevic, A. Prototypic sequences for human repetitive DNA. J Mol Evol 35, 286–91 (1992).

121. Bao, W., Kojima, K.K. & Kohany, O. Repbase Update, a database of repetitive elements in eukaryotic genomes. Mob DNA 6, 11 (2015).

122. Pagès, H., Aboyoun, P., Gentleman, R. & DebRoy, S. Biostrings: Efficient manipulation of biological strings. (2021).

123. Martin, M. Cutadapt removes adapter sequences from high-throughput sequencing reads. EMBnet.journal 17, 3 (2011).

124. Lawrence, M. et al. Software for Computing and Annotating Genomic Ranges. PLOS Computational Biology 9, e1003118 (2013).

125. Machlab, D. et al. monaLisa: an R/Bioconductor package for identifying regulatory motifs. Bioinformatics, btac102 (2022).

126. Fornes, O. et al. JASPAR 2020: update of the open-access database of transcription factor binding profiles. Nucleic Acids Research 48, D87–D92 (2020).

127. Okonechnikov, K., Conesa, A. & García-Alcalde, F. Qualimap 2: advanced multi-sample quality control for high-throughput sequencing data. Bioinformatics 32, 292–4 (2016).

128. Carroll, T.S., Liang, Z., Salama, R., Stark, R. & de Santiago, I. Impact of artifact removal on ChIP quality metrics in ChIP-seq and ChIP-exo data. Front Genet 5, 75 (2014).

129. Cox, J. et al. Andromeda: A Peptide Search Engine Integrated into the MaxQuant Environment. Journal of Proteome Research 10, 1794–1805 (2011).

130. Cox, J. et al. Accurate proteome-wide label-free quantification by delayed normalization and maximal peptide ratio extraction, termed MaxLFQ. Mol Cell Proteomics 13, 2513–26 (2014).

131. Ritchie, M.E. et al. limma powers differential expression analyses for RNA-sequencing and microarray studies. Nucleic Acids Res 43, e47 (2015).

132. Long, J.A. jtools: Analysis and Presentation of Social Scientific Data. (2020).

133. Gaujoux, R. & Seoighe, C. A flexible R package for nonnegative matrix factorization. BMC Bioinformatics 11, 367 (2010).

134. Perez-Riverol, Y. et al. The PRIDE database resources in 2022: a hub for mass spectrometry-based proteomics evidences. Nucleic Acids Res 50, D543–d552 (2022).

