## Supplementary_Information for "Readout of histone methylation by Trim24 locally restricts chromatin opening by p53"

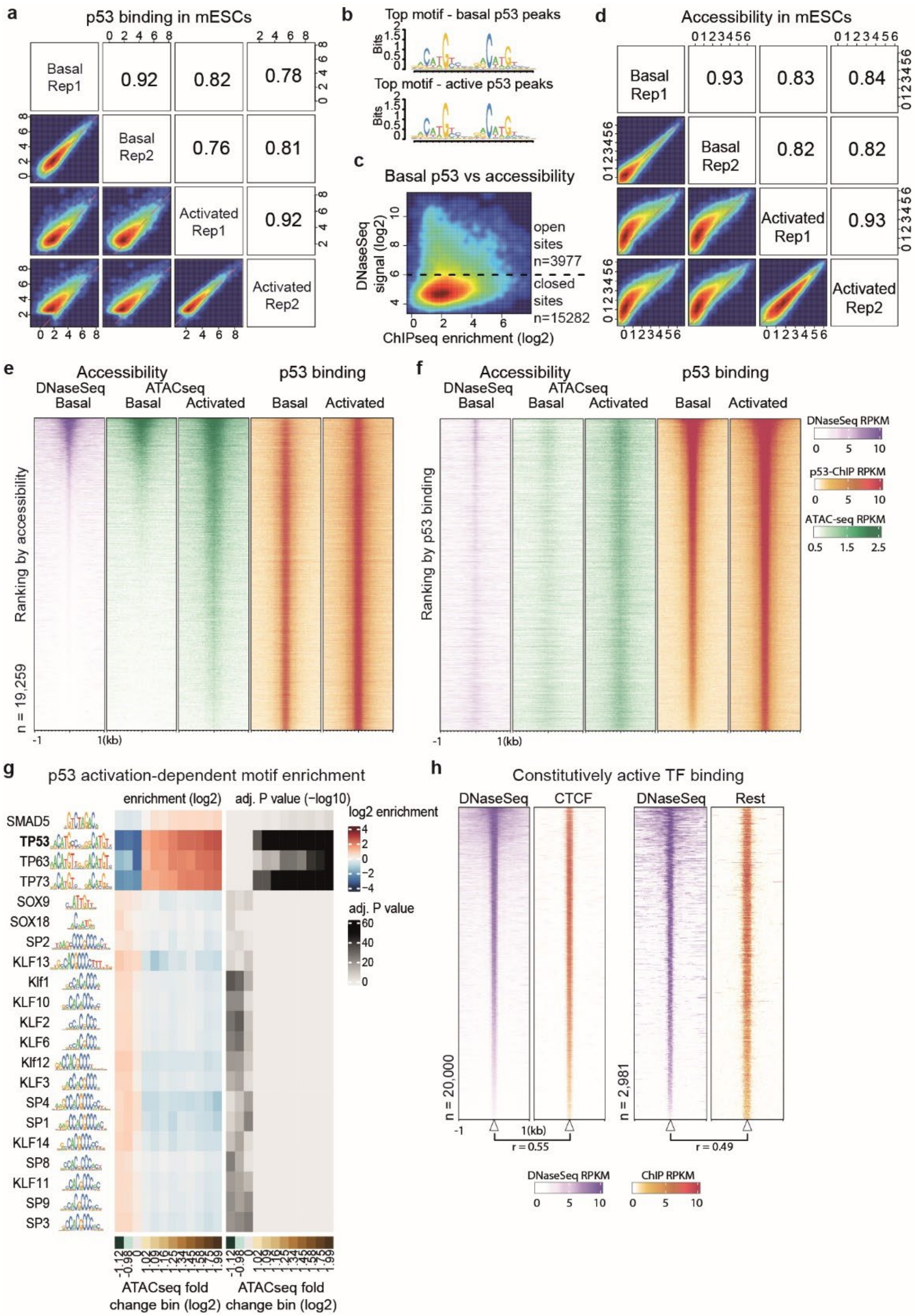


**Supplementary Figure 1 | The p53 transcription factor binds closed and open chromatin sites in mESCs and can create accessible chromatin after activation. a**) Reproducibility of p53 ChIPseq signal in mESCs, in basal and activated conditions (1uM doxorubicin, 4hrs), at the joint set of p53 peaks in both conditions (n=19259 sites). Shown are log2 enrichments over IgG control datasets for independent replicates (Rep1/Rep2). Pearson correlation coefficients are indicated. **b**) Motifs identified from the top 500 p53 ChIPseq peaks from either basal or active conditions using HOMER. **c**) p53 enrichment in basal mESCs compared to DNaseSeq accessibility signal (log2 normalized signal) in p53 peaks. Only a minority of bound sites demonstrate accessibility (> 6 log2 normalized DNaseSeq signal). **d**) Reproducibility of ATACseq signal in mESCS (log2 CPM), in basal and activated conditions, at the joint set of p53 peaks (n=19259 sites). At least half of all p53 peaks (n=9781) show substantially increased accessibility upon activation, i.e., ≥ 2-fold increase in average ATACseq signal. Pearson correlation coefficients are indicated. **e**) Heatmaps of DNaseSeq, ATACseq and p53 binding under basal and active conditions at the joint set of p53 peaks and ranked by the DNaseSeq signal. **f**) As in **e**, except sites are ranked by the average of p53 ChIPseq enrichment in basal and active conditions. RPKM values are as indicated (right). **g**) Enrichment and significance of TF motifs in mESC ATACseq peaks binned by change in accessibility upon activation of p53 (n=1500sites/bin), colour bar below indicates the minimum log2 fold change of ATACseq within each bin and centre zero fold-change bin. Shown are the top 21 enriched motifs with –log10 *P* adj. > 4 and log2 enrichment > 0.5. **h**) Heatmaps of DNaseSeq, CTCF and REST binding in mESCs as shown for the top enriched 20K CTCF sites and REST sites (n=2981), respectively. Binding of both TFs correlates well with DNaseSeq, Pearson correlation coefficients and RPKM values are as indicated (below).

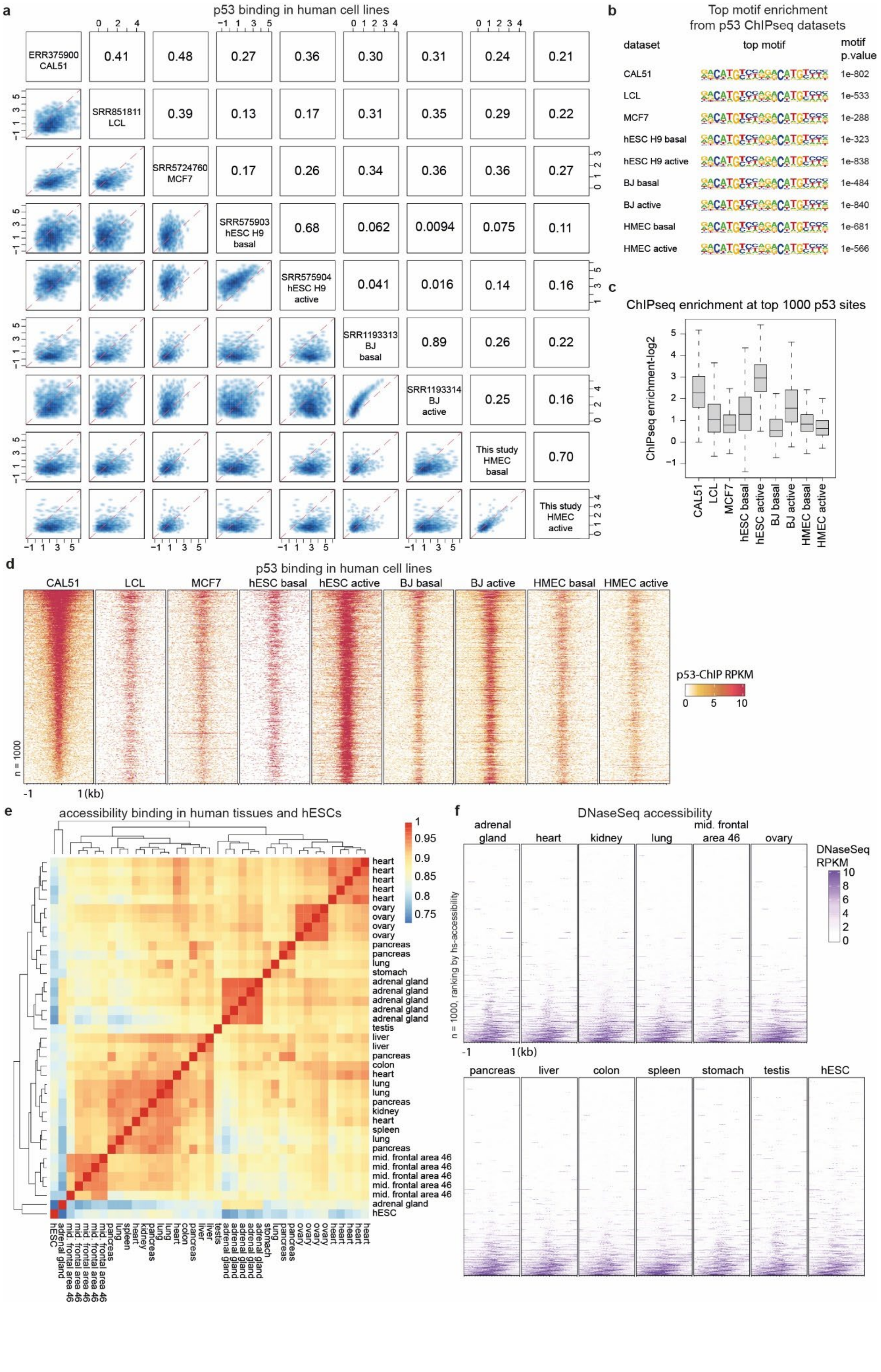
a
p53 binding in human cell lines
ERR375900 CAL51
SRR851811 LCL
SRR5724760 MCF7
SRR575903 hESC H9 basal
SRR575904 hESC H9 active
SRR1193313 BJ basal
SRR1193314 BJ active
This study HMEC basal
This study HMEC active
b
Top motif enrichment from p53 ChIPseq datasets
dataset
top motif
motif p.value
CAL51 1e-802
LCL 1e-533
MCF7 1e-288
hESC H9 basal 1e-323
hESC H9 active 1e-838
BJ basal 1e-484
BJ active 1e-840
HMEC basal 1e-681
HMEC active 1e-566
c
ChIPseq enrichment at top 1000 p53 sites
ChIPseq enrichment-log2
CAL51
LCL
MCF7
hESC basal
hESC active
BJ basal
BJ active
HMEC basal
HMEC active
d
p53 binding in human cell lines
CAL51
LCL
MCF7
hESC basal
hESC active
BJ basal
BJ active
HMEC basal
HMEC active
p53-ChIP RPKM
n = 1000
-1
1(kb)
e
accessibility binding in human tissues and hESCs
f
DNaseSeq accessibility
adrenal gland
heart
kidney
lung
mid. frontal area 46
ovary
DNaseSeq RPKM
n = 1000, ranking by hs-accessibility
pancreas
liver
colon
spleen
stomach
testis
hESC

**Supplementary Figure 2 | p53 binds sites located both at closed and open chromatin regions in human tissues and hESCs. a**) The log2 enrichments of p53 ChIPseq signal over matched control datasets in various human cell lines. Publicly available data are as indicated by dataset accession identification number, as well as cell line description and the p53-induction state as per study conditions. Data is shown at the top 1000 p53 peaks, as ranked by the mean signal across all datasets, and represent those sites that are commonly bound across diverse cell types. Pearson correlation coefficients are indicated. **b**) Top known motifs enriched from p53 binding datasets by HOMER. **c**) Boxplot showing p53 log2 ChIPseq enrichments as in **a**. **d**) Heatmaps of ChIPseq at top p53 peaks and ranked by signal across all datasets. RPKM values are as indicated (right). **e**) Correlation-based clustering of publicly available tissue and hESC DNaseSeq datasets, at the top 1000 p53 binding peaks. Tissue-source for each dataset and Pearson correlations on log2 normalized signal are as indicated. **f**) Heatmaps of DNaseSeq data from human tissues and hESCs. Averages are shown for datasets originating from the same tissue type, with RPKM values as indicated (right).

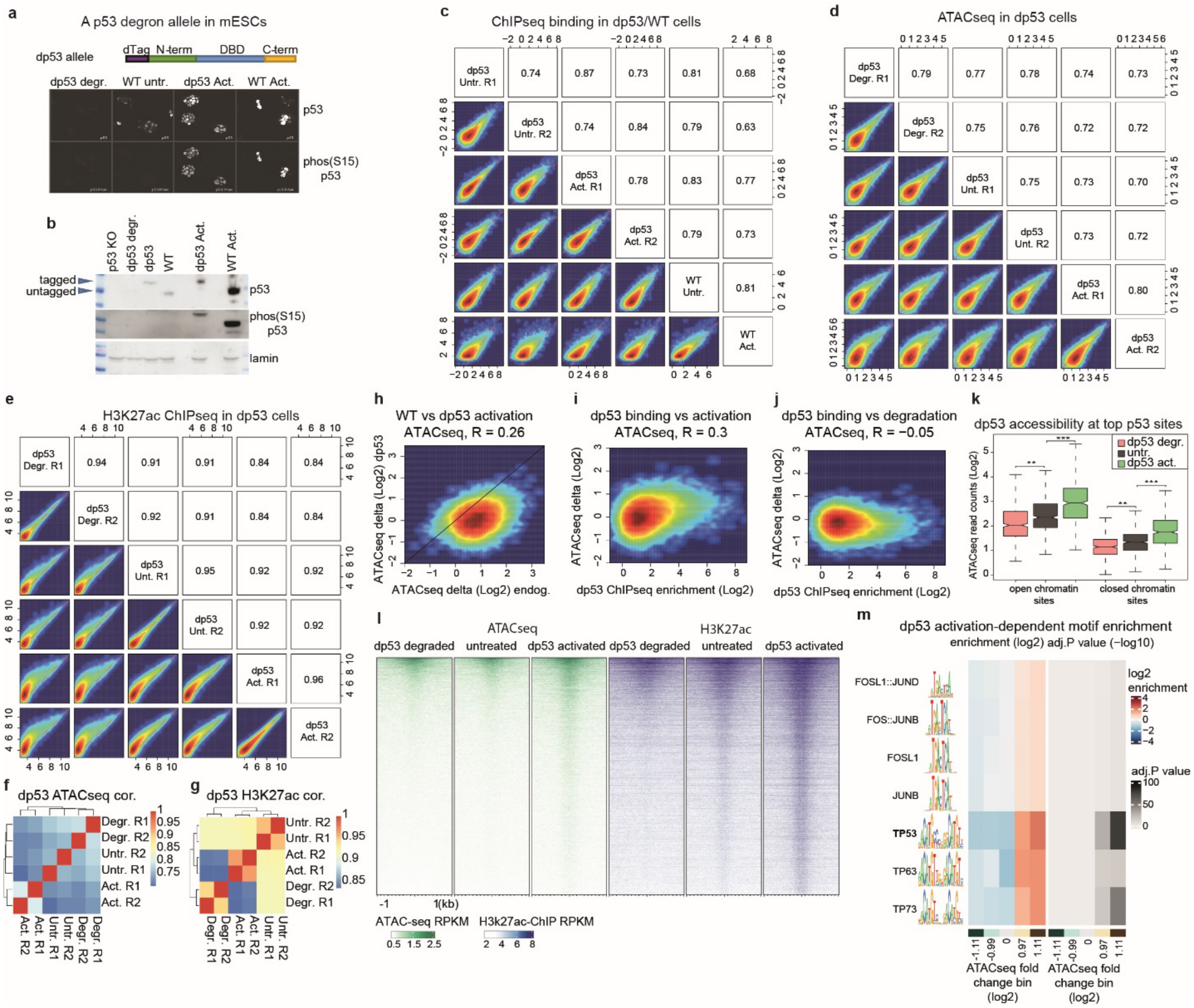


**Supplementary Figure 3 | A degron tag system for rapid removal of p53 from mESCs. a**). Schematic of *p53* with the V5 and dTag (FKBP12$^{F36V}$) sequences inserted into the endogenous gene, generating a *dp53* allele. Shown underneath are immunofluorescence visualizations of dp53 and WT p53, with either a total p53 antibody or a S15 phosphorylation p53- antibody (S18 in the mouse) specific to active p53. **b**) Western blot of p53 or p53-S15phosphorylation in *p53* knockout (KO) cells, dp53 either degraded, in basal or activated conditions and WT p53 in basal or activated conditions. Lamin serves as a loading control. Basal levels of dp53 are roughly equivalent to the WT allele and increase upon activation, though less-so than that of the activated WT allele. **c**) Reproducibility of dp53 ChIPseq signal in mESCs, in basal and activated conditions alongside average enrichment of WT p53, at the joint set of p53 peaks (n=19259 sites). Shown are log2 enrichments over IgG control datasets for replicates (R1/R2). Pearson correlation coefficients are indicated. **d**) Reproducibility of ATACseq signal (log2 CPM) in the *dp53* line, in degraded, basal and activated conditions, for replicates (R1/R2) at the joint set of p53 peaks (n=19259 sites). Pearson correlation coefficients are indicated. **e**) Reproducibility of H3K27ac ChIPseq (log2 normalized signal) in the dp53 line under degraded, basal and activated conditions for replicates (R1/R2), at the joint set of p53 peaks (n=19259 sites). Pearson correlation coefficients are indicated. **f**) Correlation-based clustering of ATACseq signal or **g**) H3K27ac ChIPseq signal in the *dp53* allele line. Pearson correlations on log2 counts are as indicated. **h**) A comparison of the log2 increase in ATACseq signal in the dp53 line and WT mESCs upon stress induction, with Pearson correlation indicated. Sites that increase in accessibility in the WT p53 line tend also to increase in the dp53 line. **i**) ChIPseq enrichment in the dp53 line against ATACseq change upon activation or **j**) degradation, compared to basal ATACseq signal. Highly bound sites tend to increase upon activation. **k**) ATACseq signal in the top 500 binding sites for the dp53 allele (n=500) that are either in open or closed chromatin in the parental mESC line (log2 DNaseSeq signal >6). *P* value ** < 0.01, *** < 0.001. **l**) Heatmaps of ATACseq and H3K27ac ChIPseq under degraded, basal and active conditions as in **d** and **e**, at the joint set of p53 binding peaks and ranked by the average signal across datasets. RPKM values are as indicated (below). **m**) Enrichment and significance of TF motifs in sites binned by change in accessibility (n=1000 sites/bin) upon activation of dp53, from all ATACseq peaks in mESCs, colour bar below indicates the minimum log2 fold change of ATACseq within each bin and centre zero fold change bin. Shown are motifs with –log10 *P*. adj > 4.

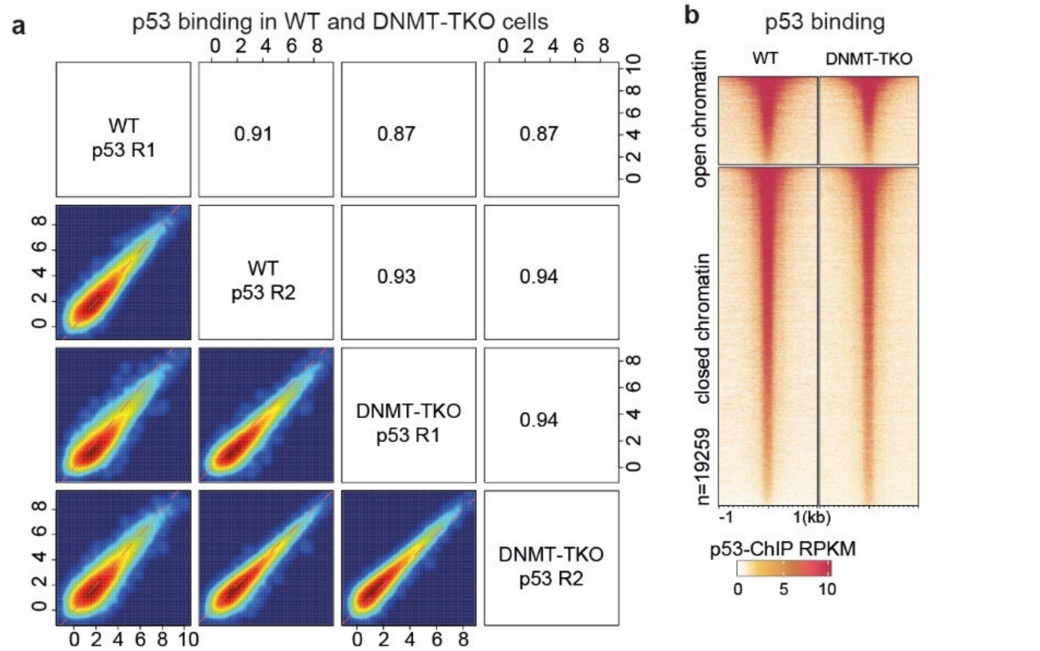


**Supplementary Figure 4 | DNA methylation does not explain diverse chromatin binding of p53 in mESCs**. **a**) Reproducibility of p53 ChIPseq signal in DNMT-TKOs, and their parental WT line, in basal conditions. Shown are log2 enrichments over IgG control datasets for replicates (R1/R2) at p53 peaks (n=19259 sites). Pearson correlation coefficients are indicated. **b**) Heatmaps of p53 ChIPseq from WT or DNMT-TKO cells at p53 peaks and separated by sites that are either in open or closed chromatin in the parental WT line (log2 DNaseSeq norm. enrichment >6). Sites are ranked by average binding signal and RPKM values are as indicated (below).

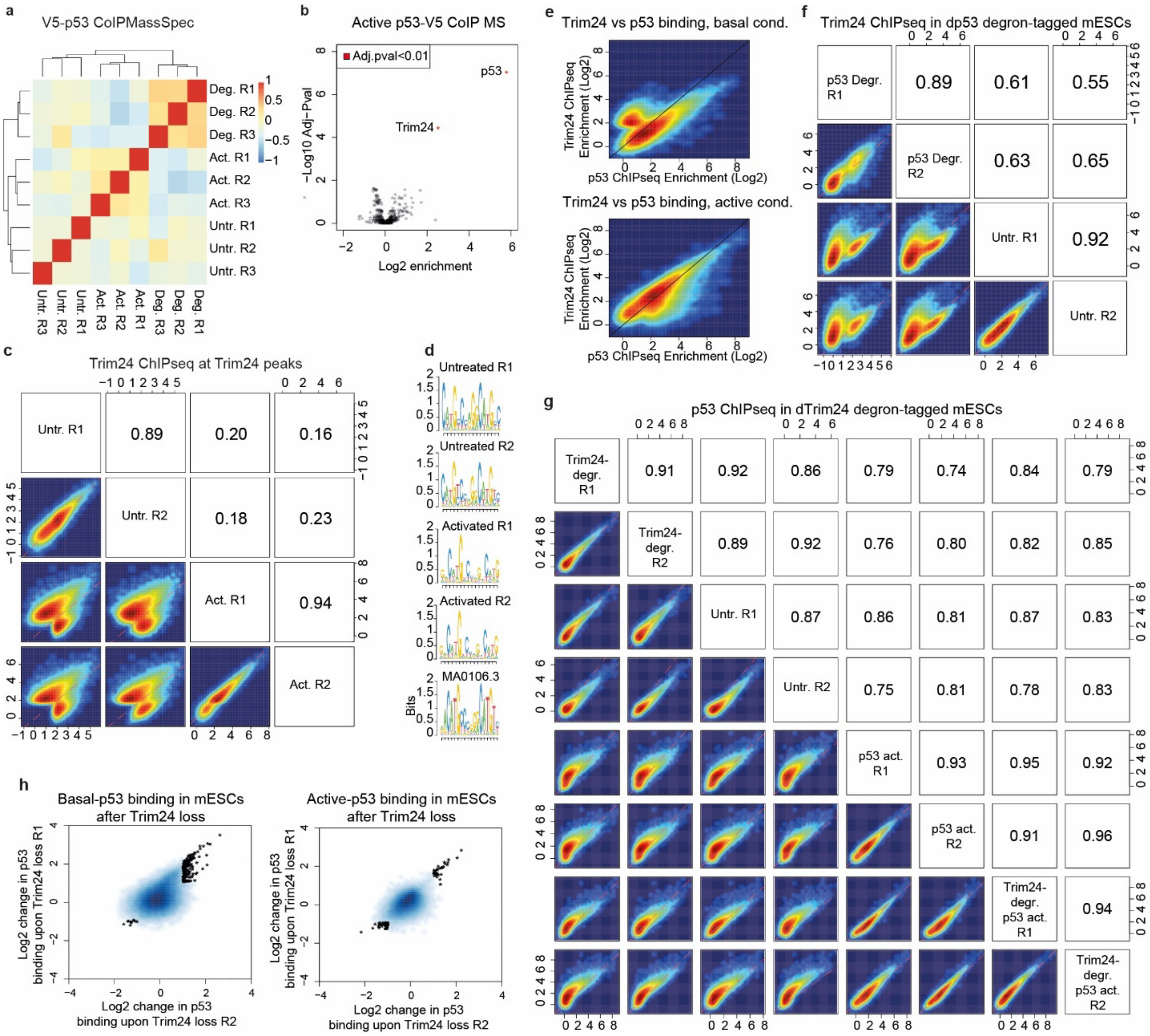


**Supplementary Figure 5 | p53 recruits Trim24 to the genome in mESCs. a**) Correlation-based clustering of mass spectrometry-identified proteins upon V5-pulldown in the dp53 degron-tagged line, under degraded, untreated or active-p53 conditions. Pearson correlations are as indicated. **b**) V5-tagged p53 CoIP enriches for both p53 and Trim24 proteins under p53-active conditions. Shown are log2 enrichments over the negative control dataset where p53 is degraded. **c**) Reproducibility of Trim24 ChIPseq signal in mESCs, in basal and p53-activated conditions, at the joint set of Trim24 peaks (n= 25768 sites). Shown are log2 enrichments over IgG control datasets for replicates (R1/R2). Pearson correlation coefficients are indicated. **d**) Top *de novo* HOMER motifs enriched from independent Trim24 ChIPseq experiments from **c**, reminiscent of the canonical p53 motif (below, Jaspar motif MA0106.3). **e**) The log2 enrichments of p53 and Trim24 at the combined set of Trim24 and p53 peaks (n= 29871), either in basal (top) or p53-active (bottom) conditions. The majority of sites are well-bound in both conditions, while a subset of p53 sites show relatively low Trim24 enrichments. **f**) Reproducibility of Trim24 ChIPseq signal in the dp53 degron-tag line, in basal and p53-degraded conditions, at all Trim24 peaks. Shown are log2 enrichments over IgG control datasets for independent replicates (R1/R2). Pearson correlation coefficients are indicated. **g**) Reproducibility of p53 ChIPseq signal in a dTrim24 degron-tag line, in basal and p53-activated conditions and upon Trim24 degradation, at the combined set of Trim24 and p53 peaks. Shown are log2 enrichments over IgG control datasets for independent replicates (R1/R2). Pearson correlation coefficients are indicated. **h**) Delta-delta plots showing the reproducibility of log2 changes in p53 binding upon degradation of Trim24. A small number of p53-bound sites shows differential enrichment (≥ 2-fold) in either basal (n=161) or active (n=69) conditions, with most increasing upon Trim24 loss.

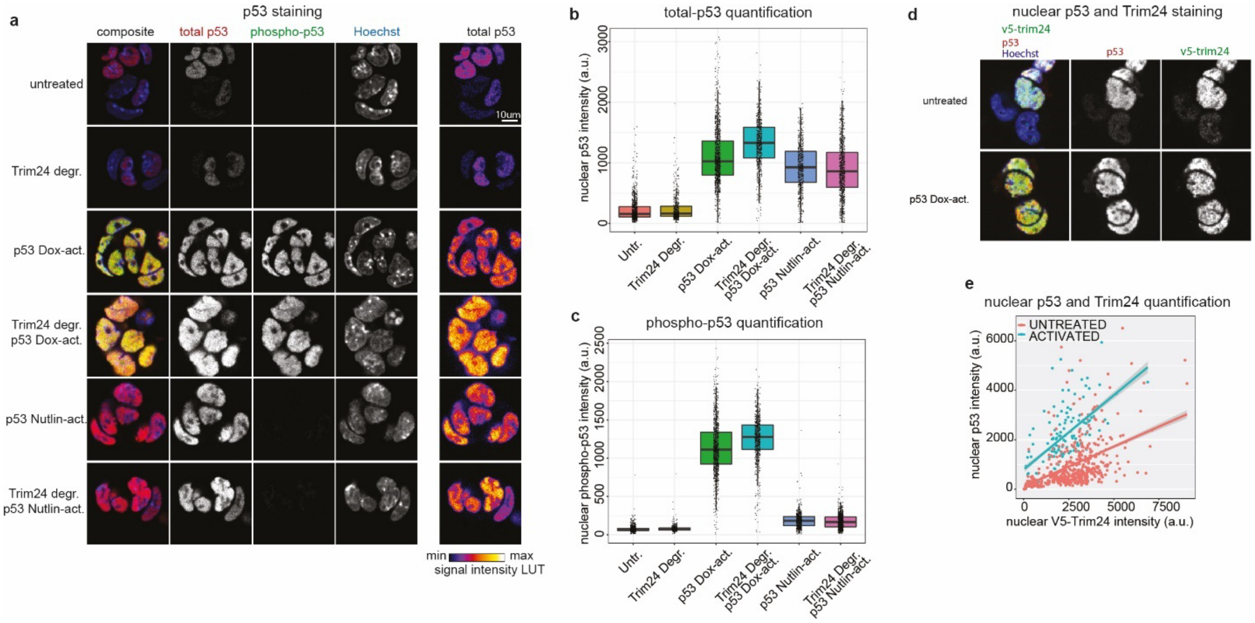


**Supplementary Figure 6 | Immunofluorescence analysis of p53 and Trim24 in mESCs. a**) Immunofluorescence visualizations of either total p53 or active S15 phosphorylation-specific p53 in the dTrim24 degron-tag line, in basal and p53-activated conditions and upon Trim24 degradation. Both doxorubicin-Dox (1µM) and nutlin3a-Nutlin (20µM) mediated activation of p53 are shown with representative images. **b**) Quantification of immunofluorescence signal in **a**, for both total and **c**) S15 phosphorylation-specific p53. a.u., arbitrary units. **d**) Visualization of either total p53 or Trim24 in the dTrim24 degron-tag line, shown are representative images. **e**) Nuclei-quantification of immunofluorescence signal in **d**, demonstrating p53 levels scale with Trim24 levels within cells, for both basal and p53-activated conditions. a.u., arbitrary units.

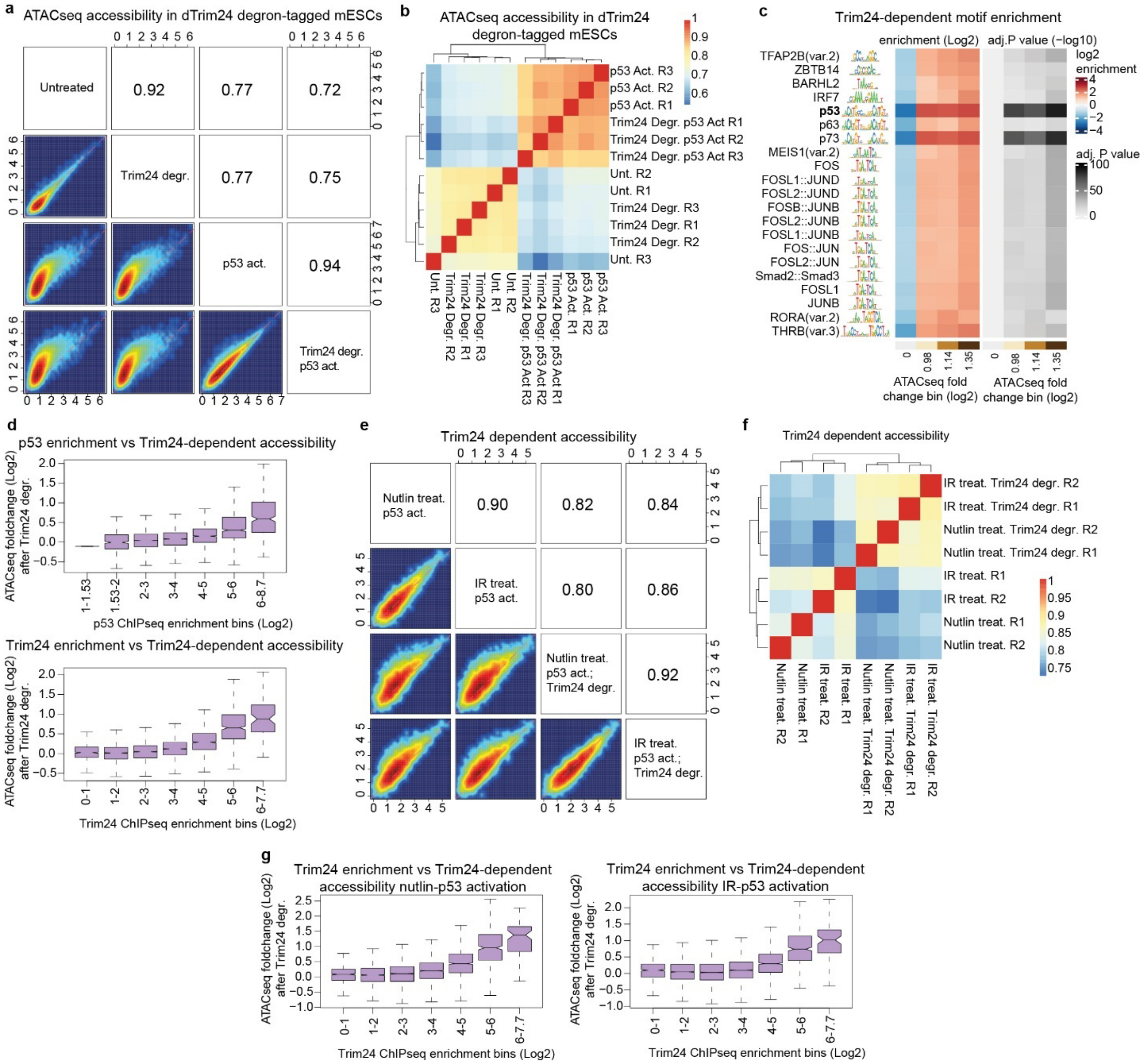

**Supplementary Figure 7 | Trim24-dependent accessibility at p53 binding sites. a**) ATACseq signal (log2 CPM) in the dTrim24 degron-tagged line, in basal and activated conditions, at the set of Trim24 ChIPseq peaks in active-p53 conditions (n=19031 sites). Shown are the average normalized signal from three replicates per condition. Pearson correlation coefficients are indicated. **b**) Correlation-based clustering of ATACseq signal in the dTrim24 degron line at the set of Trim24 peaks in active-p53 conditions. Replicates (R1/R2/R3) cluster together under p53-active conditions where Trim24-loss has an effect on p53 sites. Pearson correlations are as indicated. **c**) Enrichment and significance of TF motifs in sites binned by change in accessibility upon degradation of Trim24 in p53-active conditions (n=200sites/bin), from all mESC ATACseq peaks (n~200K), colour bar below indicates the minimum log2 fold change of ATACseq within each bin and centre zero fold change bin. Shown are the top 21 enriched motifs with –log10 *P* adj > 4. **d**) Change in ATACseq signal (log2) upon Trim24 loss in p53 active conditions, at p53-peaks and binned by increasing p53 ChIPseq enrichment (top) or Trim24 ChIPseq enrichment (bottom). **e**) ATACseq signal (log2 CPM) in the dTrim24 degron line, at the set of p53 peaks in active-p53 conditions (n=18833 sites) and upon p53 activation with either nutlin3a treatment (20uM) for 4 hours or ionizing radiation (IR) (60Gy) and left for 4 hours. Shown are the average normalized signal from two replicates per condition. Pearson correlation coefficients are indicated. **f**) Correlation-based clustering of ATACseq signal in the dTrim24 degron line upon p53 activation with either nutlin3a or irradiation and Trim24 degradation, for strong Trim24 binding sites, i.e., > 3.5-fold log2 Trim24 ChIPseq enrichment (n=3341 sites). Replicates cluster with Trim24-expression, rather than p53 activation method. Pearson correlations are as indicated. **g**) Change in ATACseq signal (log2) upon Trim24 loss and p53 activation with either nutlin3a (left) or irradiation (right) for 4 hours, at p53-peaks and binned by increasing Trim24 ChIPseq enrichment.

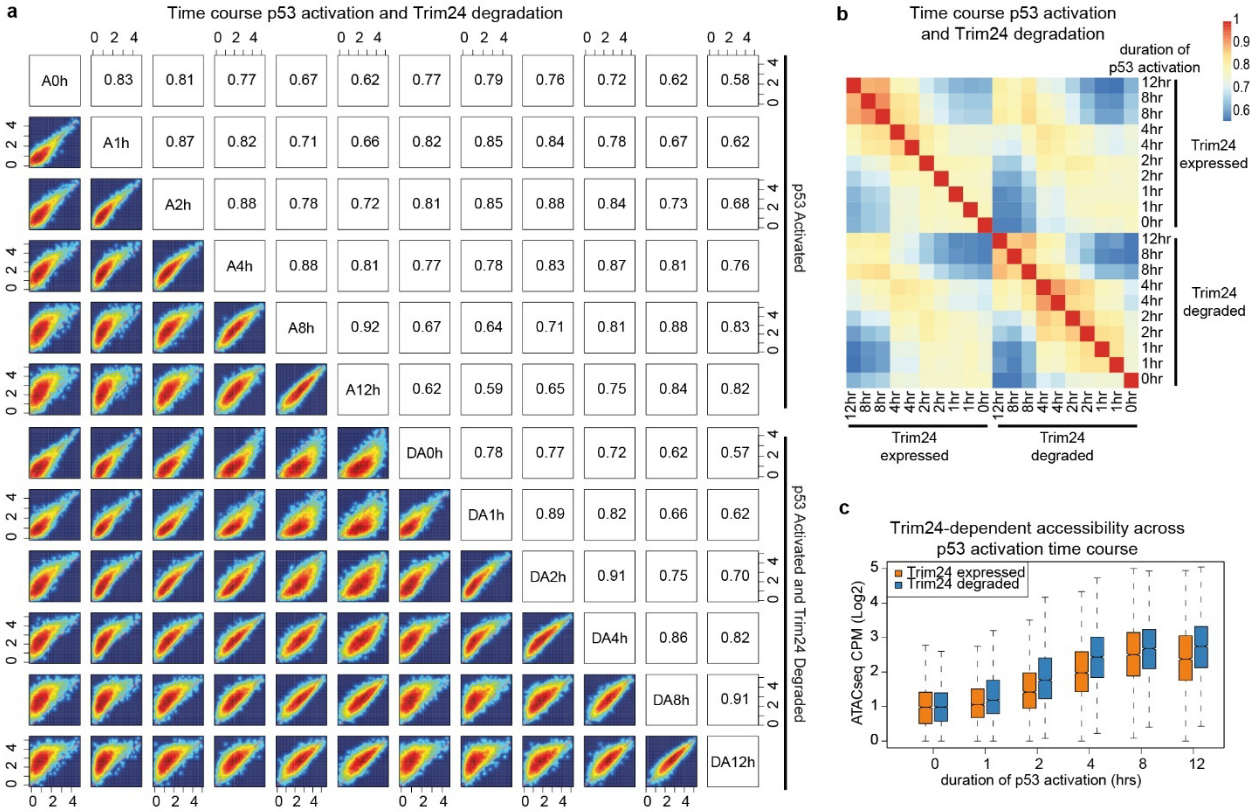


**Supplementary Figure 8 | Trim24-dependent accessibility time course. a**) Average ATACseq signal (log2 CPM) in the dTrim24 degron line, in basal and p53-activated conditions ranging from 1-12 hours after p53-activation (Doxorubicin-1µM) with Trim24 expressed (A0h-A12h) and with Trim24 degraded simultaneously (DA0h-DA12h). Shown are the set of p53 peaks strongly bound by Trim24 (log2 Trim24 ChIPseq enrichment ≥ 3.5, n=3341 sites). Pearson correlation coefficients are indicated. **b**) Pearson correlations of ATACseq signal in the dTrim24 degron line between samples upon p53 activation and Trim24 degradation in **a**. **c**) ATACseq signal at strong Trim24 peaks upon p53 activation and with or without Trim24 degradation over time. Loss of Trim24 increases accessibility, which persists up to 12 hours after p53-activation.

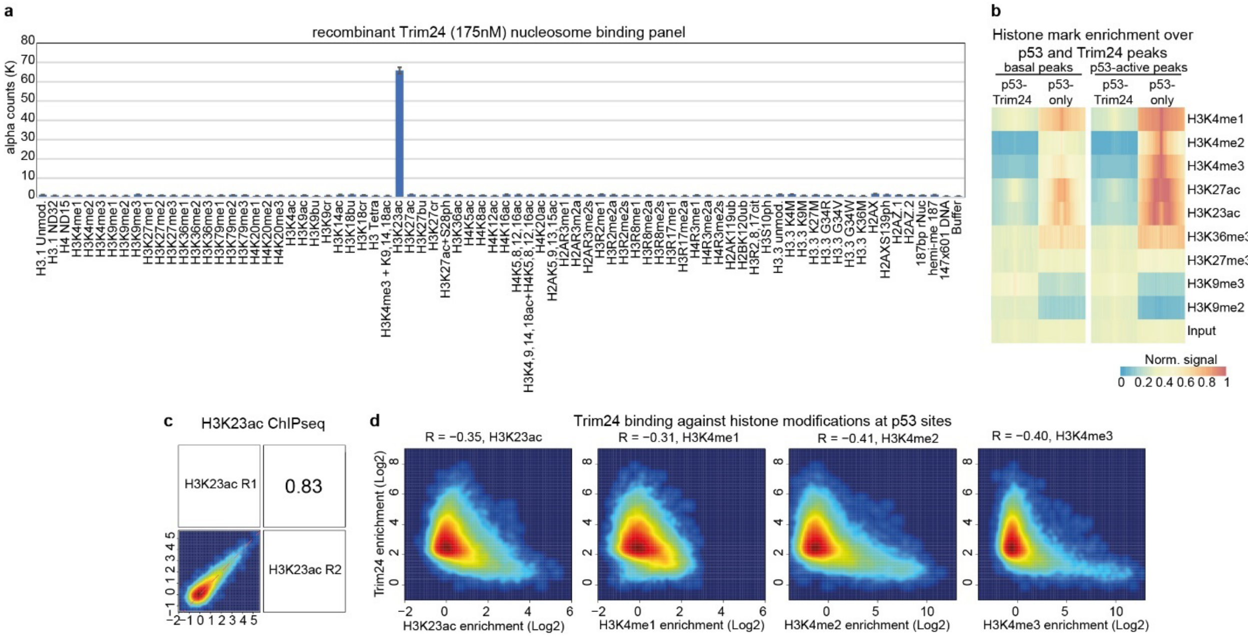


**Supplementary Figure 9 | Histone binding dynamics of Trim24. a**) Amplified luminescence proximity homogeneous assay (Alpha) binding of full-length recombinant Flag-Trim24 at 175nM against a panel of modified nucleosomes. Variant histones and/or modifications are as indicated, as well as DNA and buffer controls. Bars represent the average Alpha counts from two replicates, error bars indicate SEM. Amplified signal occurs when nucleosomes are modified with H3K23ac. **b**) Enrichment of histone modifications at Trim24-p53 co-bound peaks and p53-only peaks in basal and p53-active conditions. Shown is the normalized ChIPseq signal across 1kb regions. **c**) Reproducibility of H3K23ac ChIPseq signal in mESCs at the joint set of p53 and Trim24 peaks in p53-active conditions (n=23677 sites). Shown are log2 enrichments over IgG control datasets for independent replicates (R1/R2). Pearson correlation coefficients are indicated. **d**) Average enrichment of active histone marks, including H3K23ac and H3K4me1/2/3, against Trim24 enrichment at the joint set of p53 and Trim24 peaks in p53-active conditions. Shown are average log2 enrichments over IgG control datasets. Pearson correlation coefficients are indicated.

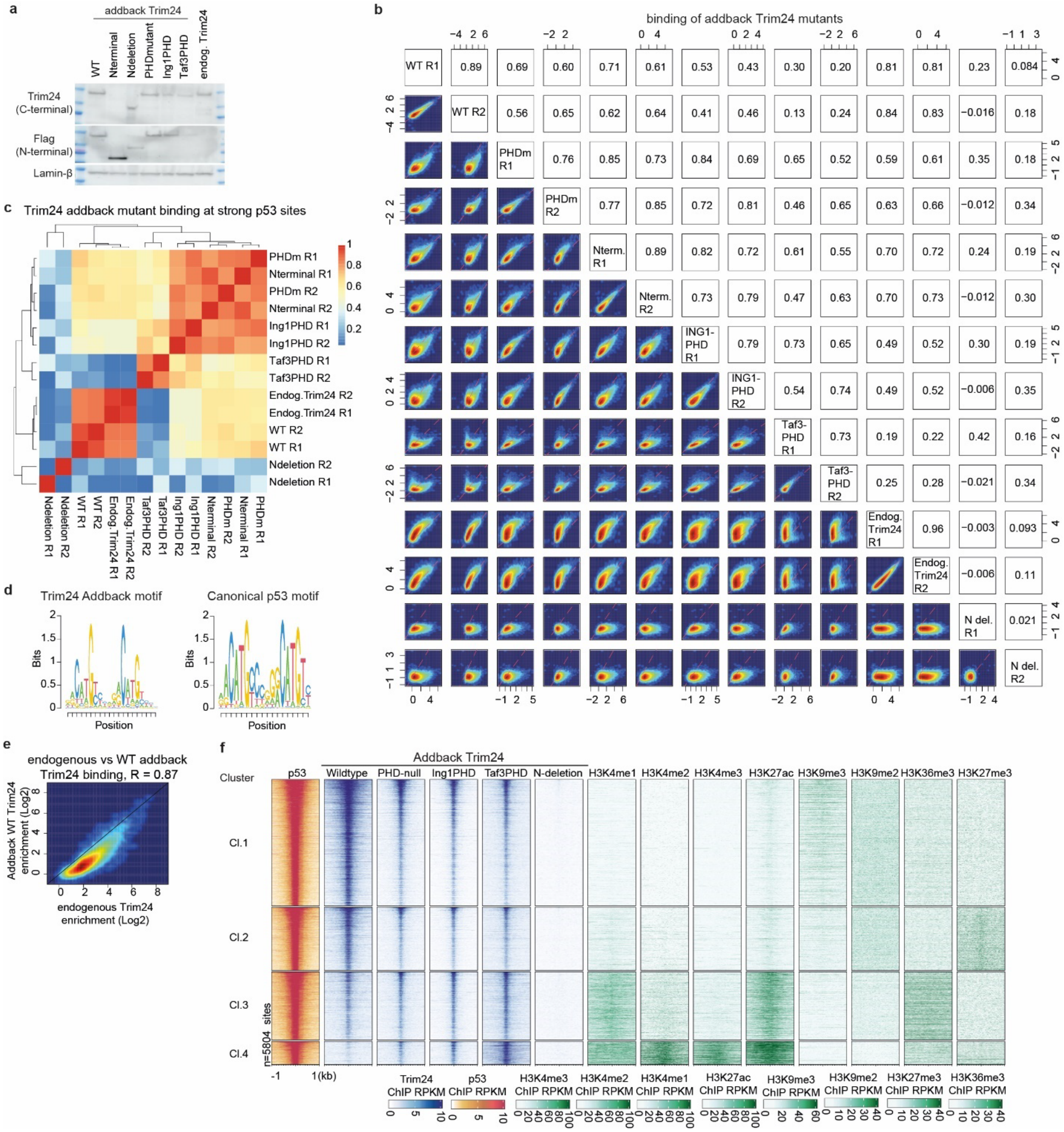

**Supplementary Figure 10 | The PHD domain of Trim24 regulates binding in closed chromatin. a**) Western blot of Flag-NLS-tagged, constitutively expressed addback-Trim24 variants in *Trim24* knockout cells, including wildtype (WT), the N-terminal fragment and N-terminal deletion of the RBBC domain, PHD-null and mutants where the PHD domain is exchanged with that of the ING1 and TAF3 protein that bind H3K4 when methylated. Addbacks are expressed at approximately endogenous levels, with the exception of the higher expressed N-terminal fragment. Lamin-β shown as loading control. **b**) Reproducibility of Flag-ChIPseq signal in addback-Trim24 expressing cells at the joint set of addback-Trim24 peaks in p53-active conditions (n= 30542 sites), as well as endogenous Trim24 ChIPseq. Shown are log2 enrichments over IgG control datasets for independent replicates (R1/R2). Pearson correlation coefficients are indicated. **c**) Correlation-based clustering of addback-Trim24 samples at strong p53 binding sites (n=5804 sites, ≥ 3.5-fold log2 p53 ChIPseq enrichment). Pearson correlations are as indicated. **d**) Top *de novo* HOMER motif enriched from addback-Trim24 peaks, strongly reminiscent of the canonical p53 motif (right, Jaspar motif MA0106.3). **e**) Average enrichment of addback-WT Trim24 against endogenous Trim24 at the joint set of endogenous Trim24/p53 peaks in active conditions. Binding decreases slightly in the addback-WT Trim24 relative to endogenous enrichments but is highly correlated. **f**) Heatmaps of ChIPseq for p53 and addback-Trim24 variants, as well as histone marks in mESCs and K-means clustered by histone marks. Sites are ranked by average enrichment across datasets. RPKM values are as indicated (below).

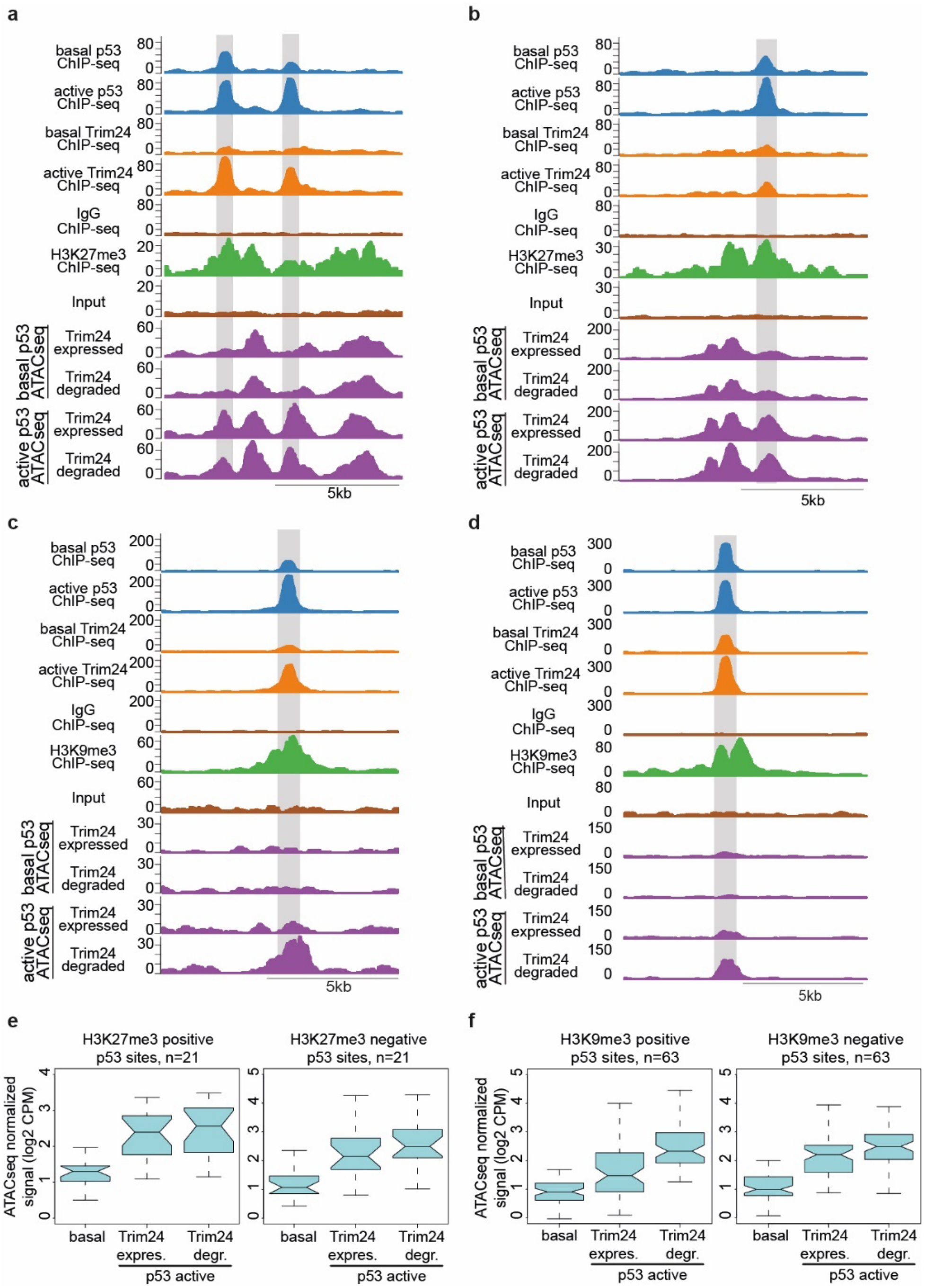


**Supplementary Figure 11 | p53 also binds and opens loci enriched for H2K27me3 and H3K9me3 histone modifications. a, b**) Representative regions of the genome where p53 and Trim24 binding (ChIPseq) colocalizes with a H3K27me3-enriched locus, as well as accessibility (ATACseq) in the dTrim24

degron line under p53 basal and active conditions and with or without Trim24-degradation. Highlighted are representative p53-Trim24 peaks. **c, d**) Representative regions of the genome where p53 and Trim24 binding (ChIPseq) colocalizes with a H3K9me3-enriched locus, as well as accessibility (ATACseq) in the dTrim24 degron line under p53 basal and active conditions and with or without Trim24-degradation. Highlighted are representative p53-Trim24 peaks. **e**) ATACseq signal at strong p53 peaks (≥ 3.5-fold log2 ChIPseq enrichment) overlapping genomic regions enriched at least 2-fold for H3K27me3 (left) or at a set of randomly sampled p53 sites (right) without H3K27me3 and with matched enrichment for p53 and H3K4 methylation levels. **f**) ATACseq signal at strong p53 peaks (≥ 3.5-fold log2 ChIPseq enrichment) overlapping genomic regions enriched at least 2-fold for H3K9me3 (left) or at a set of randomly sampled p53 sites (right) without H3K9me3 and with matched enrichment for p53 and H3K4 methylation levels.

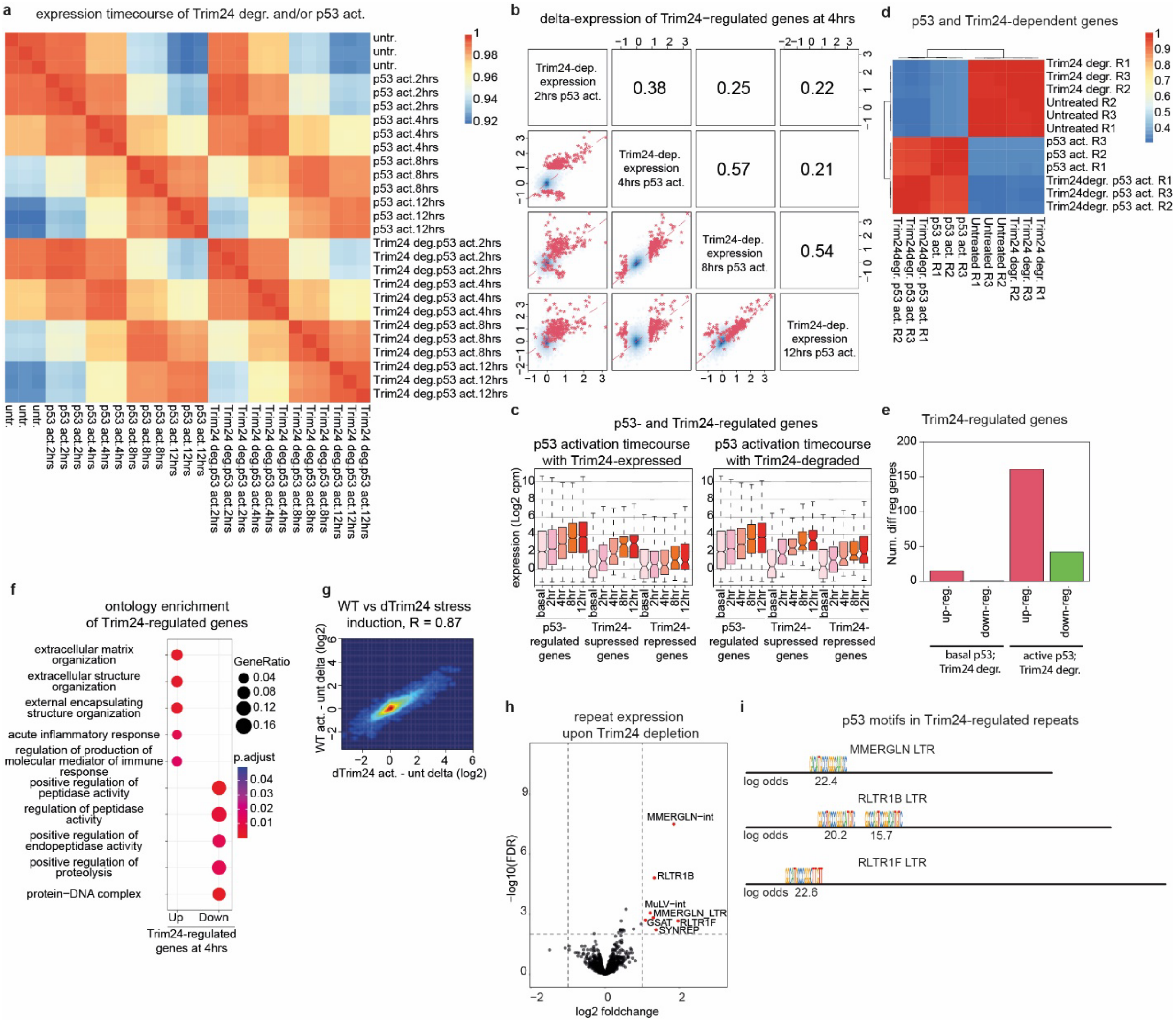


**Supplementary Figure 12 | Regulation of gene expression by Trim24 in mESCs. a**) Pearson correlations on gene expression (log2 CPM) in the dTrim24 degron line between samples upon Trim24 degradation and up to 12 hours after p53-activation (Doxorubicin-1µM), with three replicates per condition. **b**) Reproducibility of expression changes upon Trim24 degradation at various timepoints of p53-activation.

Genes that are differentially expressed at the 4hr point (shown in red, FDR<0.01 and log2 foldchange at least 0.75) show a similar response across other points in the time course of p53 activation. Pearson correlations of expression change of all genes are as indicated. **c**) Expression of p53 activated and Trim24-repressed genes (log2 CPM) at basal and different time points of p53 activation with Trim24 (left) or without Trim24 (right) in the Trim24 degron line. Shown are data from groups of differentially expressed genes at the 4-hour time point of p53 induction as in Fig. 4**a**: genes differentially expressed upon stress (p53-regulated, n=2779), genes that are differentially expressed upon stress and repressed by Trim24 (Trim24-supressed, n=100), and genes that are not differentially expressed upon stress due to repression by Trim24 (Trim24-repressed, n=103). **d**) Correlation-based clustering of RNAseq datasets in the Trim24 degron line upon loss of Trim24 in basal or p53-active conditions, for the top 500 most variable genes. Replicates (R1/R2/R3) cluster together in p53-active and basal conditions. Pearson correlations on gene expression (log2 CPM) are as indicated. **e**) Number of differentially expressed genes upon loss of Trim24 in basal or p53-active conditions (FDR<0.01 and log2 foldchange at least 0.75). Relatively few genes are differentially expressed upon loss of Trim24 in the absence of p53 activation. **f**) Gene ontology enrichment of terms associated with the Trim24-regulated genes at the 4-hour timepoint of p53 activation. **g**) Change in gene expression (log2 fold change) upon p53 activation in the Trim24 degron line compared to the WT parental line, showing highly similar effect. Pearson correlation is as indicated. **h**) Differential expression of repeat elements upon Trim24 loss in p53-active conditions at the 4-hour timepoint. Shown are the log2 fold change in RNAseq signal at repeat masker-annotated instances and grouped according to repeat subclass. Repeats that significantly increase in expression upon Trim24 loss are indicated (red). **i**) Instances of p53 motifs with a log2-odds score of at least 10 in repeat promoter elements (LTRs) of those differentially expressed repeats in **h**.

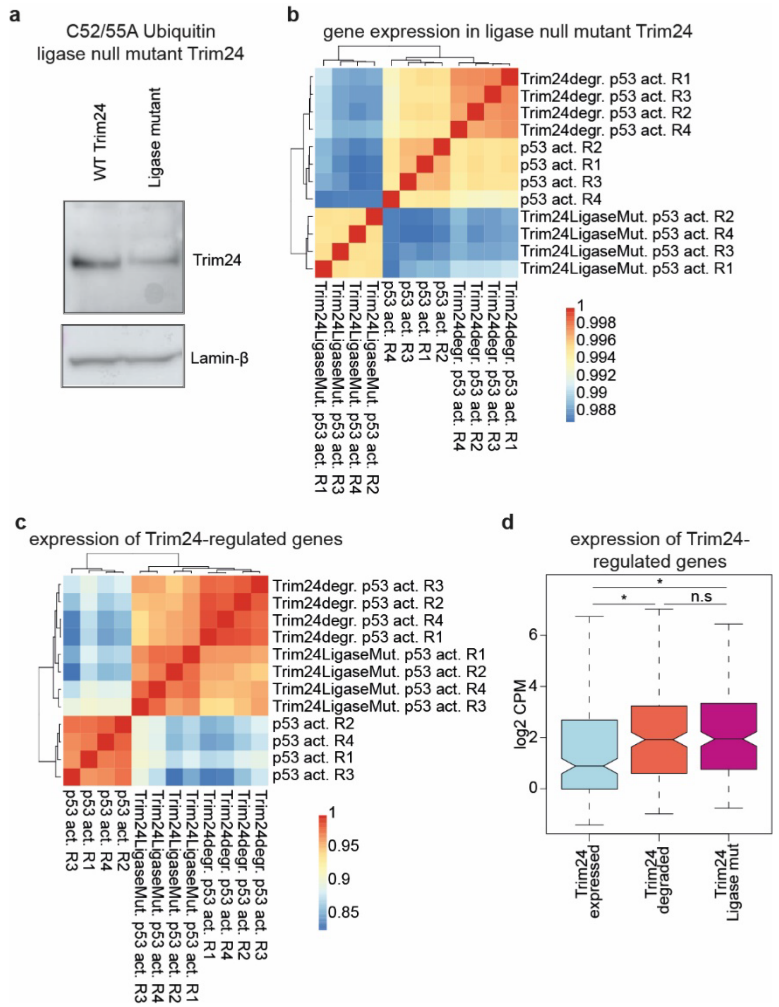


**Supplementary Figure 13 | Contribution of RING domain to Trim24 in mESCs. a**) A ligase-null variant Trim24 allele, with ammino acids mutated (C52/55A) that are essential for ubiquitin ligase activity of RING domain proteins. **b**) Correlation-based clustering of RNAseq datasets in the Trim24 degron line upon activation of p53, and the ligase-null variant Trim24 line upon activation of p53, for all genes and **c**) for Trim24-regulated genes in the degron-tagged line (n=203 genes). Ligase-null replicates (R1/R2/R3/R4) cluster together with Trim24-degraded replicates for Trim24-regulated genes. Pearson correlations on log2 CPM are as indicated. **d**) Boxplot showing expression (log2 CPM) of Trim24-regulated genes in the Trim24 degron line upon activation of p53, and the ligase-null variant Trim24 line upon activation of p53. n.s. not significant, adj. *P* value * < 0.05.

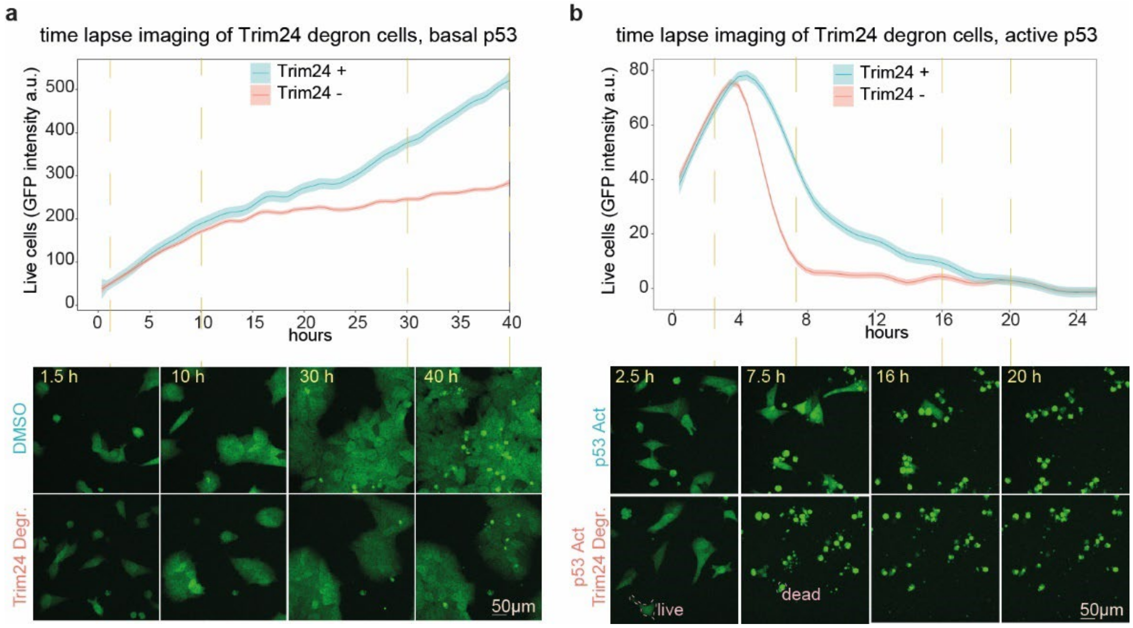


**Supplementary Figure 14 | Effect of Trim24 on cell growth and viability. a,b**) Cell growth and viability of mESCs as measured by the total intensity of fluorescence from live Trim24 degron cells expressing GFP, with and without Trim24 degradation. The average of at least 6 areas per condition is shown, shading indicates standard error around the mean. Representative images are shown at indicated points throughout the time course (below). **a**) Quantification and representative images of mESCs with or without Trim24 degradation. **b**) Quantification and representative images after stress induction (1μM Doxorubicin) and with or without Trim24 degradation. Cells initially divide but rapidly undergo apoptosis at approximately the four-hour time point. Data as in Fig. 4g, except showing time points out to 24 hours post stress induction.

**Supplementary Movie 1 |** Timelapse imaging of fluorescence from live Trim24 degron cells expressing GFP, with and without Trim24 degradation. Representative images are shown of cells either treated with DMSO (control) or dTAG compound (Trim24 Deg).

**Supplementary Movie 2 |** Timelapse imaging of fluorescence from live Trim24 degron cells expressing GFP, with p53 activated and with and without Trim24 degradation. Representative images are shown of cells either treated with doxorubicin to activate p53 (Act) or doxorubicin and dTAG compound (Act_Deg) to activate p53 and remove Trim24.
